# Profiling Gingival Inflammation in a 3D Oral Tissue Model Reveals Early Features of Disease Progression

**DOI:** 10.64898/2026.06.05.730462

**Authors:** Miryam Adelfio, Mattia Bonzanni, Grace E. Callen, Anyelo R. Diaz, Bruce J. Paster, Xuesong He, Hatice Hasturk, Chiara E. Ghezzi

## Abstract

Gingival health depends on a balanced interplay among the gingival epithelium, immune system, and oral microbiome. Disruption of this equilibrium through sustained biofilm accumulation and host inflammatory responses leads to gingivitis, a highly prevalent yet reversible condition which if left untreated could progress into more severe and irreversible condition called periodontitis. The early onset of gingivitis remains poorly defined due to subtle clinical presentation and pronounced interindividual variability. Current diagnostic approaches rely largely on clinical assessment and endpoint biomarkers, limiting insight into the early host–microbiome interactions that drive disease initiation. Here, we employ a previously validated, physiologically relevant oral tissue model (OTM) to longitudinally investigate epithelial–microbiome interactions following inoculation with patient-derived dysbiotic microbiomes from early-stage gingivitis. The OTM maintained host tissue integrity and microbial viability over a seven-day period, preserving epithelial barrier function, dynamic inflammatory responses, and disease-associated microbial signatures. Notably, we establish, for the first time in an *in vitro* platform, clinical calibration against gingival crevicular fluid (GCF), demonstrating that OTM responses recapitulate inoculum-dependent inflammatory signatures, increased microbial dissimilarity under dysbiotic conditions, and coordinated host–microbiome metabolic interactions. While pro-inflammatory responses were most pronounced at early time points, subsequent modulation toward anti-inflammatory states highlights the temporal complexity of host responses and suggests that longer culture durations may further resolve disease trajectories. Collectively, these findings validate the OTM as a robust, physiologically relevant platform that captures key features of periodontal health and inflammation. By integrating host viability, microbial ecology, and clinical benchmarking, this system enables mechanistic interrogation of early disease-driving processes and provides a translational framework for advancing predictive diagnostics and preventive therapeutic strategies in periodontal disease.

## Background

Gingivitis, an inflammatory condition of the gingiva, affects a large proportion of the global population [1,2]. It is estimated that 34.4% of adults suffer from gingivitis, with prevalence reaching up to 80% among individuals from low-income communities in the United States [3–6]. In healthy conditions, a dynamic equilibrium exists among the gingival epithelium, neutrophil-mediated immunity, and the microbiome within the gingival sulcus. This homeostasis is essential for restraining pathobiont overgrowth, preventing chronic inflammation, and maintaining both periodontal and systemic health [7–14]. However, prolonged biofilm accumulation and sustained neutrophil recruitment to the gingival sulcus drive the development of gingivitis, a reversible condition that, unlike periodontitis, a chronic inflammatory condition that affects deeper periodontal tissues, including periodontal ligament and alveolar bone, can be effectively managed via plaque removal [15–17,2]. In the absence of interventions, persistent inflammation at the epithelium may extend into the underlying connective tissue and bone, leading to periodontitis [8,13,18]. Although gingivitis does not inevitably progress to periodontitis, it is a strong risk factor; thus, restoring gingival health is crucial to prevent the gingival sulcus from becoming a reservoir of opportunistic pathogens with potential for triggering an immune response at local and systemic levels as well as dissemination of virulence factors, contributing to systemic diseases, including diabetes, cardiovascular disease, and Alzheimer’s disease [14]. The clinical heterogeneity of periodontal conditions, together with limited early diagnostic markers and tools, underscores the need for deeper mechanistic insight into host-microbiome dynamics that drive disease initiation and progression, and for the development of more effective, targeted therapeutic strategies [8,19].

Physiologically, the human body responds to an insult by activating inflammatory pathways (*e.g.,* cytokines, chemokines, and immune cell activation) [19], which manifest clinically as edema (swollen gingiva), redness, and bleeding upon stimulation with or without pain [20]. However, gingivitis presents a diagnostic challenge due to its often subtle and clinically undetectable early stages [21]. This lack of overt signs limits early detection and, therefore, early treatment, also limits the identification of clinical thresholds leading to chronic conditions [22]. Current diagnostic strategies rely on profiling inflammatory biomarkers present in the gingival crevicular fluid (GCF) and saliva, or microbial signatures in dental plaque, to establish associations between microbial composition and inflammatory status [23–25]. Furthermore, physiologically relevant experimental models are emerging to investigate disease progression through longitudinal monitoring of inflammatory responses [26,17,19]. Despite recent advances, significant limitations remain owing to the high interindividual variability in disease phenotype, the lower detectability of biomarkers compared to advanced stages of periodontitis, potential cross-contamination between salivary and epithelial exudates, and, above all, the insufficient understanding of the temporal relationship between microbiome shifts and tissue inflammation [23–26,17,19]. To address these challenges*, in vitro* tissue models provide controlled platforms to longitudinally dissect initial interactions between oral tissues and the microbiome under biologically relevant conditions. Such systems enable continuous monitoring of model responses, facilitating a more comprehensive understanding of disease initiation and progression [14]. By integrating longitudinal assessment of cytokines and chemokines release, biofilm structural dynamics, and microbial interaction networks, these models allow detailed characterization of the complex host-microbiome dialogue underlying periodontal disease. This approach may facilitate the identification of molecular fingerprints associated with the early disease stages, supporting subsequent clinical validation. Given the anatomical and metabolic complexity of the oral mucosa [14], *in vitro* models must support long-term co-culture of host tissues with patient-derived microbiomes. This capability is essential for recapitulating both homeostatic and disease-associated conditions and for enabling mechanistic investigations of periodontal inflammation. These new approach methodologies (NAMs) [27] are expected to advance existing *in vitro* strategies, which predominantly rely on simplified inflammatory conditions induced by lipopolysaccharide (LPS) [28–31] or enrichment with pro-inflammatory cytokines [32–34].

The limited availability of long-term oral tissue platforms, together with the inherent challenges of investigating disease pathogenicity using clinical and animal models [14], highlights the value of our previously developed oral tissue model (OTM) as a tractable and mechanistically informative system for investigating host–pathogen interactions [35–37]. Although periodontal homeostasis depends on coordinated interactions among the epithelial barrier, microbiome, and neutrophil-mediated immunity [38], the present work focuses on the initial epithelial-microbiome interactions and does not address downstream neutrophil-mediated pathways. To capture early inflammatory events at the epithelium interface, we investigated interactions between the gingival epithelium and patient-derived subgingival microbiomes from early-stage gingivitis within the OTM. Gingival cellular responses and microbial signatures were assessed longitudinally over seven days. To establish clinical calibration, the inflammatory patterns and microbial profiles observed in the OTM were correlated with the corresponding patient-derived GCF biomarkers and subgingival microbiome composition. Overall, our findings demonstrated that the OTM sustains interactions between host tissue and a dysbiotic microbiome for up to seven days without compromising host viability or functional responses. Moreover, responses in healthy and inflamed OTMs, clinically calibrated against GCF profiles, revealed inflammatory and metabolic host-microbiome interactions driven by microbial signatures, alongside preservation of microbiome phenotypes throughout the culture period.

Collectively, these findings expose a fundamental gap in periodontal research: the lack of physiologically relevant systems capable of resolving early host–microbiome interactions that initiate disease. Overcoming this limitation is imperative for defining early pathogenic mechanisms and for enabling the development of predictive diagnostics and preventive therapeutic strategies [39].

## Methods

### Gingival anatomical model preparation

#### Silk-based anatomical gingival scaffolds: manufacturing

Silk-based anatomical gingival scaffolds were prepared according to the established protocol, as previously described [40,37]. Briefly, *Bombyx mori* silkworm cocoons (5g) (Tajima Shoji Co., Yokohama, Japan) were cut into small pieces and degummed in 2L of deionized (D.I) water supplemented with sodium carbonate (0.02 M) (Thermo-Fisher Scientific, Waltham, MA) for 30 minutes. After degumming, silk fibers were rinsed three times in D.I. water, dried overnight, and dissolved in lithium bromide in D.I. water (9.3 M) (Sigma-Aldrich, St. Louis, MO) for 2 h at 60°C. The obtained solution was dialyzed against D.I. water for three days by means of a cellulose dialysis tube (3.5 kD MWCO, Spectrum Labs Inc., Rancho Dominguez, CA) for a total of six D.I. water changes. As a last step, silk solution was collected into tubes and centrifuged at 9000 RMP (twice) for 20 minutes at 4°C. Silk concentration (w/v) was assessed by weighing a dried sample of known volume.

To fabricate the anatomical scaffolds, silk solution (4% w/v) was pipetted into polydimethylsiloxane (PDMS) molds (̴ 4 mL/mold), recreating a three-gum-tooth unit (lower jaw) [35], degassed, and then freeze-dried (Labconco, MO) to induce formations of a porous structure using water as a porogen. Insolubility in aqueous solution was achieved by modifying the β-sheet crystalline structure of the protein via autoclave sterilization techniques, as previously reported [41]. For cell culture studies, scaffolds were autoclaved once more before seeding.

#### Oral primary gingival cells: subculture

Human primary gingival fibroblasts (hGFCs – FC-0095) and epithelial (keratinocytes) (hGECs – FC-0094) cells were purchased from Lifeline Technologies (Lifeline Cell Technologies, Frederick, MD) and cultured according to the manufacturer’s instructions. Specifically, hGFCs were subcultured in FibroLife serum-free medium supplemented with FibroLife serum-free LifeFactors (LL-0001), while hGECs in DermaLife basal medium supplemented with DermaLife K LifeFactors kit (LL-0007), as previously described [37].

#### Artificial saliva preparation

Artificial saliva was prepared as previously described [36,37]. Briefly, an antibiotic-free co-culture media was formulated by combining fibroblast basal medium with calcium chloride-depleted keratinocyte basal medium [42] (LL-0029, Lifeline Cell Technologies, Frederick, MD) in a 3:1 ratio, prior to the addition of cell growth factors, as previously described [35]. Subsequently, the co-culture media was supplemented with major salivary ionic and shear thinning constituents to mirror human saliva composition and non-Newtonian behavior [43,44] according to the following concentrations: purchased from Sigma-Aldrich, St. Louis, MO - *Xanthan gum* (0.05 wt%), ammonium nitrate (0.328 mg/mL), urea (0.198 mg/mL), lactic acid sodium salt (0.146 mg/mL), potassium citrate (0.308 mg/mL) and uric acid sodium salt (0.021 mg/mL); purchased from Thermo-Fisher Scientific (Waltham, MA) - potassium chloride (0.202 mg/mL), sodium chloride (1.594 mg/mL) and potassium phosphate (0.636 mg/mL). Complete artificial saliva solution (pH 7.4) was sterile filtered with a 100 µm strainer and a vacuum filtration system (0.45 µm) (VWR, Philadelphia).

### Inoculated Oral Tissue Model (OTM)

#### Native shear stress stimulation: bioreactor fabrication

The long-term culture bioreactor was manufactured at the Lawrence Lin MarkerSpace-UMass Lowell, as previously described [36]. The housing consisted of a main chamber equipped with a lid made in Derlin® acetal resin (McMaster-Carr, Elmhurst, IL) and sealed with pin screws, flat washers, wing nuts, and an O-ring (McMaster-Carr, Elmhurst, IL). A peristaltic pump (EZO-PMPTM-Atlas Scientific, Long Island City, NY) with relative tubing and connectors McMaster-Carr, Robbinsville, NJ; United States Plastic Corporation, Lima, OH; Masterflex, Vernon Hills, IL and Atlas Scientific, Long Island City, NY) was used for flow stimulation. An Arduino set-up (board, breadboard and code (https://atlas-scientific.com/peristaltic/ezo-pmp/) was adopted to control input and output signals. Pumps were calibrated daily to ensure flow rate accuracy. Prior to experiments, all components were autoclaved at 121 °C for 20 minutes.

#### Silk-based anatomical gingival scaffold: human gingival cell seeding

Cell populations were seeded (**Figure 1**) in the anatomical scaffold, as previously described [35]. Briefly, gingival scaffolds were equilibrated in phosphate buffer saline (PBS 1X). Subsequently, hGFCs were detached from tissue culture flasks and embedded in a neutralized solution of rat tail type I collagen before being seeded onto the open porosity of the scaffold at a density of 200,000 cells/mL. Neutralized collagen solution was prepared in ice by incorporating acidic type I collagen (First Link, UK) and Dulbecco’s modified Eagle medium (DMEM, 10x - Sigma-Aldrich, St. Louis, MO) in a 4:1 ratio and neutralized with 10M sodium hydroxide (Sigma-Aldrich, St. Louis, MO). During collagen polymerization at 37°C, hGECs were gently retrieved from culture flasks and seeded twice, onto the closed porosity of the scaffold at a seeding density of 50,000 cells/cm^2^ with an interval of 2 hours between seedings. Cellularized gingival scaffolds were cultured submerged in antibiotic-free co-culture medium for two days and subsequently in antibiotic-free artificial saliva throughout the culture period (2 weeks). On day 5, gingival scaffolds were grown in air-liquid interface (ALI) configuration to induce hGECs stratification and differentiation. Artificial saliva changes were performed every other day until day 7 and then every day after the microbiome inoculum. For nomenclature purposes, tissue maturation of the fully cellularized models (hereafter referred to as mature anatomical models) was achieved during the initial seven days of culture. Host–microbiome interactions were subsequently evaluated over a separate seven-day period following microbial inoculation, designated as Day 0 (microbiome inoculation into the mature anatomical model) through Day 7 throughout the manuscript.

**Figure 1:**
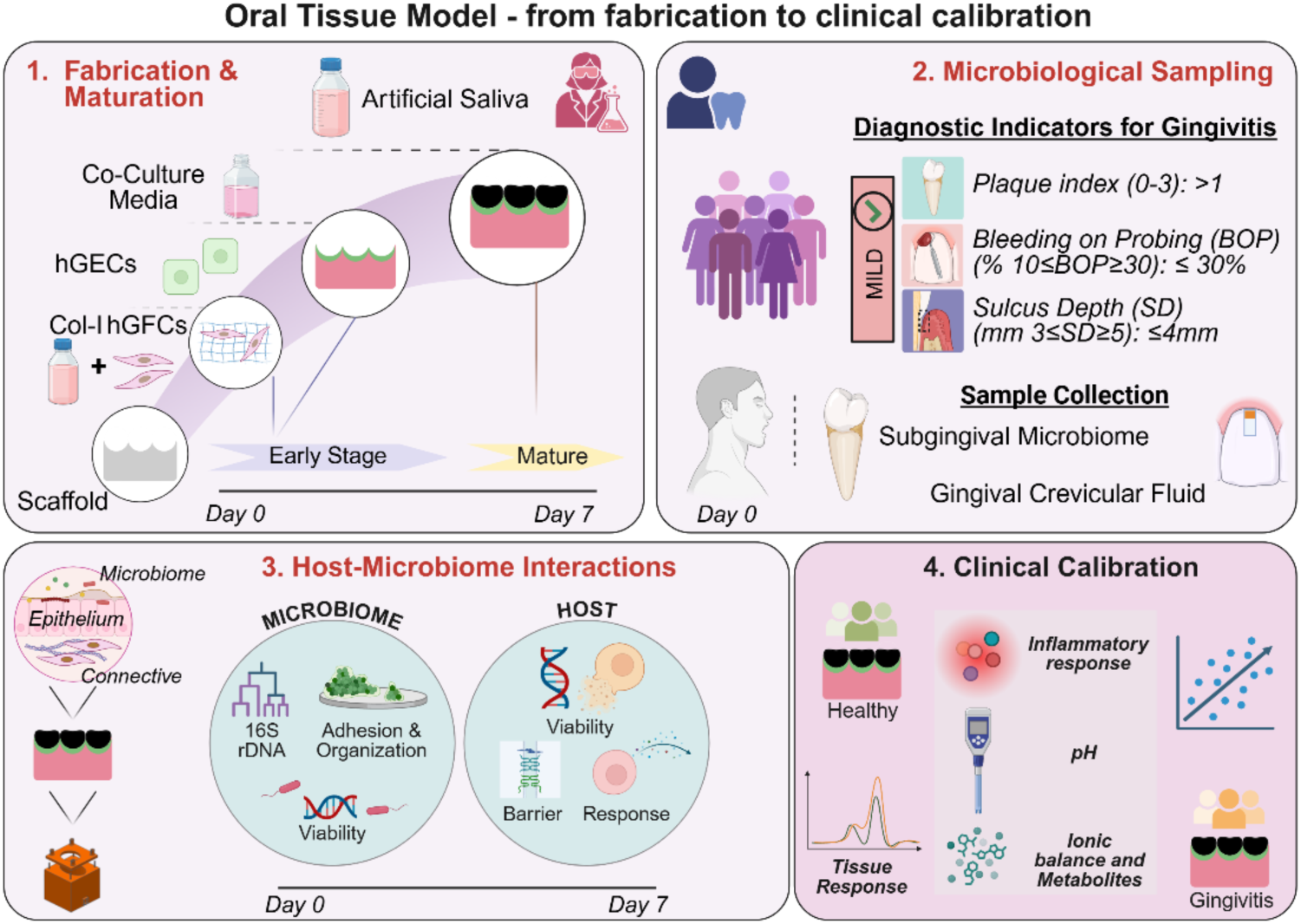
Gingivitis experimental conditions and clinical calibration of an Oral Tissue Model (OTM): experimental workflow. 1) Anatomical gingival tissue fabrication and maturation (Day 0 – Day 7) — silk-based anatomical scaffolds were populated with primary human gingival fibroblasts (hGFCs) in collagen type I, followed by primary human keratinocytes cells (hGECs). Gingival models were grown in optimized co-culture media before being cultured in artificial saliva until maturation (first seven days), referred to in the manuscript as a mature anatomical model. Host–microbiome interactions were assessed over a separate Day 0–Day 7 period. 2) Microbiological sampling from human subjects (Day 0) — Gingival Crevicular Fluid and subgingival microbiomes were sampled from subjects affected with gingival inflammation (gingivitis) upon diagnostic indicators. 3) Long-term host-microbiome interactions (Day 0 - Day 7) — upon inoculum, OTMs were cultured in native regime with a previously validated gingival bioreactor; host-microbiome interactions were longitudinally assessed by molecular and functional readouts. 4) Clinical calibration — the longitudinal performance of OTMs exposed to microbiomes from healthy [37] and gingivitis-affected subjects was compared to the pro/anti-inflammatory response relative to the gingival crevicular fluid and the pH, ionic balance, and metabolites. Cartoon was made in Biorender.com.

#### Cellular anatomical gingival scaffold: oral subgingival plaque microbiome inoculum

Collection of human subgingival plaque microbiome samples (**Figure 1**) from gingivitis donors (n=20 with 10 sampling sites/donor) was conducted at the Center for Clinical and Translational Research at the ADA Forsyth Institute, Somerville (MA) (**Supplementary Table 1** – enrollment plan) according to the published methods [45,46] and as previously described [37]. Briefly, a trained dentist assessed gingival inflammation (mild inflammation) according to diagnostic indicators: plaque index (PI), bleeding on probing (BoP) and sulcus depth (SD) [20,47–49]. Clinical presentation of mild inflammation was correlated with a PI>1, BOP≤30% and SD≤4mm. Subsequently, the surfaces of 10 separate subgingival sites/subjects were scaled by a trained hygienist using sterile Gracey curettes and plaque was preserved in Tris–Ethylenediaminetetraacetic (EDTA) buffer medium (0.5 mL- 1 tube/human subject). Plaque samples (de-identified and coded) were inoculated into the gingival pocket shortly after collection.

Before microbiome inoculation in the OTM (mature anatomical models), plaque samples were pooled. Aliquots from the initial inoculum were collected for baseline analysis of 16S rDNA analysis (*h*plaque0). 10 µL aliquots (5 µL per pocket side) were inoculated into the gingival sulcus of each scaffold. Inoculated OTMs (n=5) were cultured in static conditions at ALI for 16h (Day 0 – Day 1) to favor microbiome attachment and organization before being cultured in native regime for 7 days (salivary flow at 1 mL/minute) (Day 1 to Day 7). All the conducted studies were performed in compliance with the approved protocols: FIRB# 18-06, ADA Forsyth Institutional Review Board and IRB—20-090, the University of Massachusetts Lowell. Prior to specimen collections, all subjects signed the FIRB-approved informed consent.

#### Salivary shear stimulation (native regime)

The *ad hoc* bioreactor system delivered salivary shear stimulation at the gingival sulcus. The day after inoculation, OTMs were transferred into the bioreactor chamber and cultured in shear stimulation (native regime - 1 mL/minute) at 37°C and 5% CO_2_, with physiologically relevant shear stress parameters previously validated [36]. Longitudinal host-microbiome interactions were assessed every 24h (**Figure 1**) by sampling spent media aliquots from the chamber reservoir and from all three periodontal pockets before proceeding with specific molecular and functional assessments.

#### Oral hygiene in the OTM

Inoculated OTMs underwent oral hygiene rinses using Listerine® mouthwash (brown color, no ethanol) on Day 3 and Day 5 [37]. Briefly, a 1% v/v mouthwash solution was prepared in PBS (1X) and sterile filtered (0.2 µm, VWR, Philadelphia) before being poured into a glass beaker. Subsequently, the OTMs were transferred to the beaker, with mouthwash volume approaching ALI, and left to soak for 10 seconds. OTMs were briefly rinsed with PBS (1X) and gently dabbed on gauze before resuming the native regime.

### Host-Microbiome interactions in the OTM – Host’s assessments

#### Cell viability: lactate dehydrogenase (LDH) and picoGreen

Host’s viability was assessed via LDH (proxy of cytotoxicity) and picoGreen (DNA quantification) assays. LDH assay (n=5) was carried out according to the manufacturer’s instructions (Sigma-Aldrich, St. Louis, MO). At each time-point, 500 µL of spent saliva collected from the tissue culture well or from the chamber reservoir were stored at −80°C, before being diluted 1:5 in LDH assay buffer. The obtained LDH data was interpolated using a standard curve.

DNA aliquots (n=4) was quantified using Quant-iT picoGreen dsDNA assay kit (Thermo-Fisher Scientific, Waltham, MA), according to the manufacturer’s instructions. To ensure quantification of host cells only, cells were separated from microbial population using Percoll^TM^ (Cytiva, Marlborough, MA) density gradient [50], as previously described [37].

Briefly, the gingival epithelium and the attached microbial population were isolated from the OTM via nasal swabs and recovered by vortexing and centrifugation (500 x*g*, 5 minutes). Supernatants containing the bacterial cells were collected for viability testing (day 7) (see section: *Microbiome viability: confocal laser scanner microscopy (CLSM) and flow cytometry*). Pellet containing the epithelial cells was resuspended in 1 mL of isolation medium composed of 950 µL of 50% v/v Percoll^TM^ in PBS (1x) and 50 µL of DermaLife basal medium supplemented with 10% Fetal Bovine Serum (Thermo-Fisher Scientific, Waltham, MA), L-glutamine (6 mM) (Lifeline Cell Technologies, Frederick, MD), and 2.5 nM of L-cysteine (Thermo-Fisher Scientific, Waltham, MA). The resulting solution was centrifuged at 11,000 x *g* for 15 minutes, then the first 500 µL containing only the epithelial cells were mixed with PBS (1X) and centrifuged again at 500 x *g* for 5 minutes. The remaining 500 µL containing the microbiome population was combined with the bacterial supernatant previously collected. To ensure that no bacterial DNA was processed, the recovered epithelial cell suspension was resuspended in PBS (1X) and subsequently filtered through a 12-µm hydrophilic polycarbonate Nucleopore (Whatman) filter. To confirm cell recovery and quantify epithelial density, the membrane was briefly vortexed, and cells were lysed in 0.05% Triton-X (Sigma-Aldrich, St. Louis, MO) at −80°C before proceeding with a standard picoGreen protocol.

#### Gingival epithelial barrier: TransEpithelial Electrical Resistance (TEER)

Integrity and functionality of the gingival barrier were assessed using a volt-ohm-meter (EVOM2) and the associated 6 mm cell culture cup chamber (ENDOHM-6G) (World Precision Instrument, Sarasota, FL). On Day 7, OTMs (n=6) were punched into 7 mm discs (VWR, Philadelphia), transferred to a Transwell insert (#29442-082, VWR, Philadelphia) and TEER was recorded. Due to differences in disc thickness, TEER values were reported in Ω/mm, with thickness measured using a caliper. Background from acellular anatomical scaffolds was used to calculate net TEER [37].

#### Oral mucosal response: human β-defensing-2 anti-microbial peptide

Release of anti-microbial peptide human β-defensin 2 (hBD2) (PeproTech, Cranbury, NJ—900-K172) in OTMs was quantified via Enzyme-Linked Immunosorbent Assay (ELISA) [37]. To perform the assay, reagents and consumables were purchased separately: from Sigma-Aldrich, St. Louis, MO - Tween-20, Bovine Serum Albumin, and ABST liquid substrate solution; from Thermo-Fisher Scientific, Waltham, MA – PBS (10X); from Corning, NY – 96-well microplates. For each time-point (Day 0 −1 – 2 – 3 – 5 - 7), 100 µL of epithelium exudates (spent-artificial saliva) were collected from each periodontal pocket (n=5), briefly spun down to remove any bacterial or debris interference, and stored at −80°C until analyzed. hBD2 assay was carried out according to the manufacturer’s instructions, with samples diluted 1:4 in reagent diluent. Optical Density (OD – 405 nm) was determined by means of a microplate reader (SpectraMax M2 with SoftMax Pro7 software, Molecular device, San Jose, CA) with wavelength correction set at 650 nm. All OD values were interpolated based on a standard curve.

#### Oral mucosal inflammatory response: Milliplex® assay

Pro- and anti-inflammatory responses were conducted via Milliplex® assay [37]. Briefly, epithelial exudates (n=5) were collected on Day 0, Day 1, Day 3 and Day 7 from the gingival sulcus and processed for simultaneous analysis of multiple chemokine and cytokine biomarkers via magnetic bead panel (MilliPlex MAP, Millipore, Sigma): Granulocyte-macrophage colony-stimulating factor (GM-CSF), Interleukin 1 alpha (IL-1α), Interleukin 1 beta (IL-1β), Interleukin 1 Receptor Antagonist (IL-1RA), Interleukin 2 (IL-2), Interleukin 6 (IL-6), Interleukin 8 (IL-8), Interleukin 10 (IL-10), Interleukin 12p40 (IL-12p40), Interleukin 17A (IL-17A), Monocyte Chemoattractant Protein 1 (MCP-1), Tumor Necrosis Factor-alpha (TNF-α). Data collection was performed using the Luminex 200 instrument (Luminex, TX), with the median fluorescence intensity (MFI) in each sample recorded using the xPonent software (Luminex, TX). Raw data values were then interpolated relative to each standard curve via the Belysa® Immunoassay curve fitting software (Merck Millipore, Burlington, MA). Protein concentration (pg) was normalized on cellular density obtained by Percoll^TM^ (see section “*Cell viability: lactate dehydrogenase (LDH) and picoGreen*”) and rescaled with min-max normalization.

### Host-Microbiome interactions in the OTM – Microbiome assessments

#### Microbiome viability: confocal laser scanner microscopy (CLSM) and flow cytometry

Microbiome viability and distribution in the OTM (n=2) were qualitatively and quantitatively assessed via CLSM and flow cytometry, respectively [37]. Briefly, prior to inoculum in the OTM, an aliquot of the pooled subgingival plaque microbiome was pre-stained with Syto9 (1:250) (Thermo-Fisher Scientific, Waltham, MA) in PBS (1X) for 20 minutes at room temperature (RT). After Syto9 binding to microbial nucleic acids, the microbiome was inoculated into the OTM as described above. To assess overall viability (live/dead), on Day 7, the gingival pockets were incubated with propidium iodide (PI, 30 µM) (Thermo-Fisher Scientific, Waltham, MA) in PBS (1X) for 20 minutes at RT, followed by a quick rinse in PBS (1X) to remove excessive binding. Regionality in the OTMs (upper, middle, and lower regions) along the z-axis of the gingival sulcus was imaged with a Leica SP8 CLSM (Leica, Germany) to visualize microbiome distribution and overall viability.

Flow cytometry viability assessments were conducted on OTMs on Day 7 (n=2). Microbial samples were collected as described above (see section *“host-microbiome interactions - Cell viability: lactate dehydrogenase (LDH) and picoGreen*) and processed for live/dead assay (**Supplementary Figure 2a**) according to the published protocol [37]. In brief, the recovered microbial samples were resuspended in 0.85% sodium chloride buffer (Thermo Fisher Scientific, Waltham, MA) and divided into two main groups (5 controls): live controls (unstained, stained with Syto9 or Syto9+PI) and dead controls (stained with PI or Syto9+PI). To induce death, samples were incubated at RT for 30 minutes in 70% isopropanol with shaking intervals. Live/Dead controls were incubated with Syto9 (20 µM) and/or PI (60 µM) or left unstained for 15 minutes at RT in 0.85% sodium chloride buffer, before being fixed in 2% paraformaldehyde (PFA) (Thermo Fisher Scientific, Waltham, MA) in PBS (1X). To quantify viability, microbial population were analyzed using an Amnis® FlowSight® Imaging Flow Cytometer (Millipore Sigma, Burlington, MA), collecting 10,000 events per sample. A custom gating strategy was applied to select single-cell subpopulations (**Supplementary Figure 2a**). Compensation matrices were generated from subgingival microbiome samples prior to acquisition [51] and data analysis was carried out using the IDEAS software (Amnis Corporation, Seattle, WA). Out-of-focus events were excluded using the Gradient RMS (raw mean square) function on the free-field channel. Cells (single events) were then identified based on area *versus* aspect ratio to remove debris. The resulting population was gated for Syto9 and PI to determine the percentage of dye-positive cells.

#### Microbiome spatial organization within the gingival sulcus: scanning electron microscope (SEM)

To characterize microbial populations’ attachment and spatial distribution in the gingival sulcus, OTMs (Day 7, n=2) were briefly rinsed with PBS (1X) and then fixed in PFA 4% (Thermo-Fisher Scientific, Waltham, MA) in PBS (1X) for 2 h at RT. To image the depth of sulcus, OTMs were cut longitudinally and sliced transversally, rinsed in D.I. water, and subsequently dehydrated through a graded ethanol series (in %: 50,70,80,90,95,100) (Thermo-Fisher Scientific, Waltham, MA) followed by Critical Point Dryer treatment (Tousimis Samdri, Rockville, MD). Before imaging, OTMs were coated with Au/Pd particles (Cressington 108) (Cressington Scientific Instruments, UK) and imaged at 10 kV accelerating voltage and low vacuum (60 Pa) using a Phenom XL G2 Desktop (Thermo-Fisher Scientific, Waltham, MA) equipped with an EDS sensor.

#### Operational Taxonomic Units (OTUs) assessments: gDNA isolation

Genomic DNA (gDNA) isolation of subgingival microbiome on Day 0 (*h*plaque0) and on Day 7 (native d7) was carried out to assess microbiome preservation within the OTM in gingivitis conditions via 16S rDNA sequencing according to the published pipeline [37]. On Day 0 (*h*plaque0), 10 µL of the pooled subgingival plaque microbiome (n=5) were combined with 390 µL of PBS (1X) and pre-treated with propidium monoazide (PMA, 50 µM) in order to amplify only the microbial gDNA from live bacteria. PMA incubation consisted of 10 minutes at RT on a shaker covered with aluminum foil, followed by 15 minutes of photolysis with a blue LED device (Biotium, San Francisco). On Day 7 (native d7), gDNA isolation was performed by manually dividing one pocket/OTM (n=5) into upper, medium, and lower regions, using a razor blade, according to oxygen content [35]. Each region was transferred to a 1.5 mL tube and dissected using dissection scissors (VWR, Philadelphia) before being resuspended in 400 µL of PBS (1X). Subsequently, PMA was added to each OTM region during the pre-treatment phase with a total photolysis incubation time of 30 minutes. Post-PMA treatment, samples were centrifuged for 10 minutes at 5,000 x*g* and processed for gDNA isolation (Lucigen, Middleton, WI, USA). To induce bacterial lysis, microbial populations were resuspended in Tris-EDTA (TE) buffer supplemented with Ready-Lyse Lysozyme Solution and incubated overnight at 37°C. gDNA extraction was performed according to the manufacturer’s instructions, with total gDNA content quantified using a nanodrop (Thermo-Fisher Scientific, Waltham, MA). 300 ng of gDNA/sample were processed for sequencing analysis, as indicated by Zymo Research’s guidelines (Irvine, CA).

#### Operational Taxonomic Units (OTUs) assessments: library preparation and 16S rDNA sequencing

Library preparation and 16S rDNA sequencing were performed by Zymo Research (Irvine, CA). Using Quick-16S™ NGS Library Prep Kit (Zymo Research, Irvine, CA), gDNA samples were prepared for targeted sequencing. To ensure the best coverage of the 16S gene and high sensitivity, Zymo Research (Zymo Research, Irvine, CA) designed primers (Quick-16S™ Primer Set V1-V3). Amplicons were quantified by qPCR fluorescence and pooled at equimolar concentrations. To ensure high quality, the final pooled library was purified using Select-a-Size DNA Clean & Concentrator™ (Zymo Research, Irvine, CA), then quantified with TapeStation® (Agilent Technologies, Santa Clara, CA) and Qubit® (Thermo Fisher Scientific, Waltham, WA). Sequencing of the final libraries was performed on Illumina® MiSeq™ with a V3 reagent kit (600 cycles), with 10% PhiX spike-in.

#### Operational Taxonomic Units (OTUs) assessments: data analysis

A comprehensive data analysis and interpretation of the reads was carried out at the ADA Forsyth Oral Microbiome Core (FOMC, Somerville, MA) in accordance with the following pipeline [52]. Amplicon read processing was performed using the DADA2 pipeline, which models and corrects sequencing errors generated by Illumina platforms. The workflow consisted of five steps: (1) quality-based read trimming, (2) estimation of error rates, (3) inference of amplicon sequence variants (ASVs), (4) merging of paired-end reads, and (5) chimera removal. The resulting ASVs were subsequently used for microbial taxonomic assignments. To determine whether the identified taxa are specific to the oral microbiome, each sequence was cross-referenced against several reference databases: Expanded Human Oral Microbiome Database (eHOMD v15.2) [53], HOMD 16S rRNA RefSeq extended v1.1, Mouse Oral Microbiome Database (MOMD v0.1) [54], Greengene Gold (GG), and the NCBI 16S rRNA reference sequence database; collectively these databases comprise 25,120 sequences representing 15,601 oral and non-oral microbial species [55]. To match each OTU to species-level assignments, an alignment-based algorithm was used with NCBI BLASTN v2.7.1+ [56,57], with parameters consisting of sequence identity thresholds ≥98% and alignment length ≥90% relative to reference sequences [55]. When the reads matched multiple species with identical percentages and lengths, potential chimeras were evaluated using USEARCH v8.1.1861 [58]. This alignment-based approach allows for robust species-level taxonomic classification while maximizing species-level resolution. The read counts, calculated as 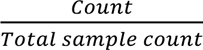 for each sample, were used to calculate taxonomic abundance [59–62].

Venn diagrams were used to quantify shared and unique OTUs between the initial inoculum (*h*plaque0) and Day 7 samples. The relative abundance data were binarized (0:absent; 1:present), and species were counted to identify shared and unique taxa. Species found to be absent from both groups were excluded from the analysis [37].

Diversity (richness, evenness, and similarity) indices between the initial inoculum (*h*plaque0) and the microbial population at Day 7 in the OTM (native d7) were assessed via α- and β-diversity indices. Differences in richness and evenness between groups were calculated by means of α-diversity (Observed features, Shannon, and Simpson) indices. Statistical differences were calculated using the Kruskal–Wallis *H*-test (alpha-group-significance function QIIME 2 “diversity” package). Similarity comparisons between *h*plaque0 and Day 7 were assessed using the β-diversity index by means of principal coordinates analysis (PCoA). Given the compositional nature of microbiome data, counts were subjected to centered log-ratio (CLR) transformation. Euclidean distances calculated using the CLR transformation (Aitchison distance) were visualized on PCoA plots. Statistical significance was assessed using PERMANOVA (permutational multivariate analysis of variance) (beta-group-significance function QIIME 2 “diversity” package).

Differential abundant taxa were identified using Linear Discriminant Analysis Effect Size (LEfSe), which considers the compositional nature of the microbiome data when comparing the proportion of different species in a sample. Differences between microbial communities were calculated by applying a rank-based Kruskal–Wallis sum-rank test between the groups of interest to identify taxa with significantly different relative abundance. Subsequently, an unpaired Wilcoxon rank-sum test was computed to determine biological consistency [63]. Effect sizes via linear discriminant analysis (LDA) scores were determined using significant differential abundant taxa.

The social structure of the microbiome was evaluated by generating a similarity matrix (https://software.broadinstitute.org/morpheus/) from species relative abundance values based on Spearman rank correlation. The resulting data was imported into a custom-made MATLAB (MathWorks Inc., MA) script [64]. Specifically, the community_louvain.mat script was applied to compute modularity and assign each species to a module, with the following parameters: similarity matrix; gamma = 1 (classic modularity); ‘negative_sym’ for symmetric treatments of negative weights. The assigned module information was used to re-arrange the order of the species in the heatmap, highlighting the modular structure of the microbial populations.

### Predictive pipeline for clinical calibration in the OTM based on epithelial exudates

#### Gingival Crevicular Fluid (GCF) sample processing

To evaluate alignment of the baseline inflammatory status of the different enrolled subjects’ tissues against the OTM’s state after microbiome inoculation, epithelial exudates (GCF) from gingivitis subjects (n=19) were collected prior to microbiome sampling by extraction with PerioPaper® strips placed for 30 seconds at the gingival sulcus (10 collection sites/subject) and subsequently snap-frozen before storing them at - 80°C. GCF extraction was carried out at the ADA Forsyth Multiplex Core (FOMC, Somerville, MA). To detect pro- and anti-inflammatory cytokines, strips were eluted with cold assay buffer (MilliPlex MAP, Millipore, Sigma) in 1.5 mL tubes and shaken for 30 minutes at 4°C. Subsequently, each strip was inserted into a spin basket (Spin-X™ centrifuge tube insert #7200120, Thermo-Fisher Scientific, Waltham, MA) connected to the same 1.5 mL tube containing the previous eluate and centrifuged for 10 minutes at 9,600 x*g* at 4°C. Aliquots from the extracted exudates were diluted 1:2 in L-AB buffer (MilliPlex MAP, Millipore, Sigma) prior to conducting the Milliplex® assay, as described above. In order to perform comprehensive clinical assessments according to disease states (healthy *vs.* gingivitis), GCF from healthy subjects (n=19) was extracted following the same procedure. Analytes assessed were: Granulocyte-macrophage colony-stimulating factor (GM-CSF), Interleukin 1 alpha (IL-1α), Interleukin 1 beta (IL-1β), Interleukin 1 Receptor Antagonist (IL-1RA), Interleukin 2 (IL-2), Interleukin 3 (IL-3), Interleukin 6 (IL-6), Interleukin 8 (IL-8), Interleukin 10 (IL-10), Interleukin 12p40 (IL-12p40), Interleukin 17A (IL-17A), Monocyte Chemoattractant Protein 1 (MCP-1), tumor necrosis factor-alpha (TNF-α). Additionally, GCF was screened for hBD2 expression, as described above.

#### Selection of biomarkers for clinical calibration based on GCF

Both Multivariate Analysis of Variance (MANOVA) and Linear Discriminant Analysis (LDA) assume that the variables being analyzed are not too highly correlated with one another (multicollinearity). Excessive correlation makes the estimates unstable and prevents the reliable determination of each variable’s unique contribution to group separation. To address multicollinearity, Pearson’s correlation coefficients were calculated for all GCF cytokines (MATLAB, MathWorks Inc., MA). In pairs with a correlation coefficient greater than 0.75, one cytokine of the pair was excluded based on biological relevance. By removing one cytokine from each pair, the analysis ensures that the remaining biomarkers meet the required assumptions of MANOVA and LDA, producing stable test statistics and interpretable discriminant function coefficients for classifying subjects. MANOVA was then performed (SPSS) to assess overall differences in cytokine profiles between the healthy and gingivitis groups. LDA (SPSS) was carried out after confirming significant (p<0.05) MANOVA results. Standardized canonical discriminant function coefficients were extracted, normalized relative to their sum, and ranked in descending order of magnitude to determine cytokine importance in the discrimination of the two groups. Posterior probabilities of group membership were calculated for each subject to evaluate the classification accuracy of the cytokines.

#### Profiling of inflammatory and microbial signatures in the OTM according to disease states

To evaluate OTM inflammatory state based on the initial inoculum (healthy *vs* gingivitis), longitudinal profiles of IL-1α, IL-1β, IL-8, and MCP-1 (selected based on normalized weight ranking (%)) were analyzed in both conditions. The release of antimicrobial peptide (hBD2) was also analyzed in relation to clinical conditions. Microbial signatures in the OTM between healthy (*h*plaque0 *vs.* native d7) and gingivitis (*h*plaque0 *vs.* native d7) were analyzed via 16S rDNA sequencing. Comparisons between samples were evaluated based on relative abundances, shared and unique OTUs (Venn analysis) and diversity (α and β) indices, as previously described.

#### Profiling ionic and metabolic response in the OTM according to disease states

To assess whether local biochemical milieu favors gingival inflammation over health, we interpreted ionic and metabolic changes at the gingival sulcus in the OTM in both periodontal health and gingivitis. Ionic and metabolic response in the OTMs (n=5/disease states) were carried out using an automated cell culture analyzer, the BioProfile FLEX2 (Nova Biomedical, Waltham, MA, USA), based on: carbon metabolism (glucose, lactate, glutamine and glutamate), ionic extracellular environment (sodium, potassium, and calcium), and nitrogen metabolism (ammonium) [67–69]. Briefly, epithelial exudates (200 µL/pocket) from each OTM were collected daily (Day 0 through Day 7), briefly spun to remove cellular debris, and stored at −80°C until analysis. Prior to quantification, samples were thawed on ice and gently mixed to ensure homogeneity. To ensure measurements were within the validated analytical range, samples were diluted 1:4 in artificial saliva. Metabolites concentrations (ι1millimole/L; ι1g/L) were calculated by subtracting values from artificial saliva (background). Positive deviations from baseline were interpreted as net production, whereas negative deviations were interpreted as net consumption.

#### Oxygen tension and pH at the gingival sulcus

To validate the distribution of the microbiome within the gingival sulcus while monitoring local metabolic changes, oxygen and pH were measured within the pocket. Oxygen measurements over time (n=5) were recorded using a PreSens’s Profiling Oxygen Microsensor PM-PSt7 equipped with a Microx 4 fiber optic oxygen transmitter (PreSens, Regensburg, Germany) on Day 0 (mature anatomical models) and Day 1. Recordings were carried out by guiding the optical fiber through the gingival sulcus (z-axis, mm) using the M-152 three-axis direct transmission micromanipulator and the GJ-1 magnetic holder with a steel base plate from Narishige (Narishige, Amityville, New York). pH (n=5) was longitudinally (Day 0 – Day 7) monitored at the gingival sulcus using the Orion™ 9810BN Micro pH Electrode (Thermo-Fisher Scientific, Waltham, MA) connected to the FiveEasy™ pH/mV Meter (Mettler-Toledo, Columbus, OH).

### Statistical analysis

Statistical significance was carried out using OriginPro 2026 (OriginLab Corporation, Northampton, MA, USA). Statistical significance (p<0.05) among normally distributed data was computed using paired T-Test, Two-sample Student’s *T*-test (two-tailed), Two-sample Student’s *T*-test (two-tailed) with Welch correction for unequal variance, One-Way ANOVA, One-Way or Two-Way repeated measures ANOVA with Bonferroni and Dunnet’s post-hoc. Statistical significance assessed through MANOVA for clinical calibration was computed in IBM SPSS Statistics. 16S rDNA sequencing data analysis was performed by ADA Forsyth Oral Microbiome Core (FOMC) at the ADA Forsyth Institute (Somerville, MA) applying Kruskal–Wallis *H*-test and PERMANOVA (QIIME 2 “diversity” package), and Kruskal–Wallis sum-rank test and unpaired Wilcoxon rank-sum test (LEfSE), with a *p*<0.05.

## Results

### OTM long-term response to dysbiotic microbiomes

In our previous manuscript [37], we demonstrated that OTM mirrored eubiotic interactions between host and healthy microbiomes, preserving both viability and stability of the human cell populations, the topography of native tissues and replicating healthy physiological responses. In the present study, we extend this platform to investigate long-term interactions between host tissue and dysbiotic microbiomes derived from subjects affected by gingivitis (**Figures 1-2**). Viability assessments (**Figure 2a**) confirmed sustained cell viability throughout the culture period. Notably, the initial increase in LDH release observed following microbiome inoculation was followed by a steady decline, indicating stabilization of cytotoxic responses over time (**Supplementary Figure 1a**). These findings were consistent with reduced epithelial density in the absence of significant tissue damage. After seven days of culture, epithelial barrier integrity remained preserved (**Figure 2b**) and comparable to OTM response in healthy conditions (**Supplementary Figure 1b**). Initial and longitudinal OTM responses to microbial challenges were evaluated by quantifying the release of anti-microbial peptide (hBD2) and pro- and anti-inflammatory cytokines (**Figure 2c-d**). Normalized hBD2 (**Figure 2c**) increased after inoculation (Day 0 – Day 1) and remained elevated throughout the culture period (Day 1 to Day 7). Hierarchical clustering further resolved the temporal dynamics of inflammatory cytokine response within the OTM (**Figure 2d)**, identifying four distinct clusters. Cluster 1 was highly expressed at Day 0, showed moderate release (IL-8, IL-1Rα, and GM-CSF) or decline (MCP-1, IL-12p40, IL-6) at Day 1, and was undetectable at later time points. Cluster 2 exhibited low-level expression (except for TNF-α) at Day 0, peaked at Day 1, and declined thereafter, with IL-1α remaining detectable. Clusters 3 and 4 were absent at early time points (Days 0 and 1) but increased at Days 3 and 7.

**Figure 2:**
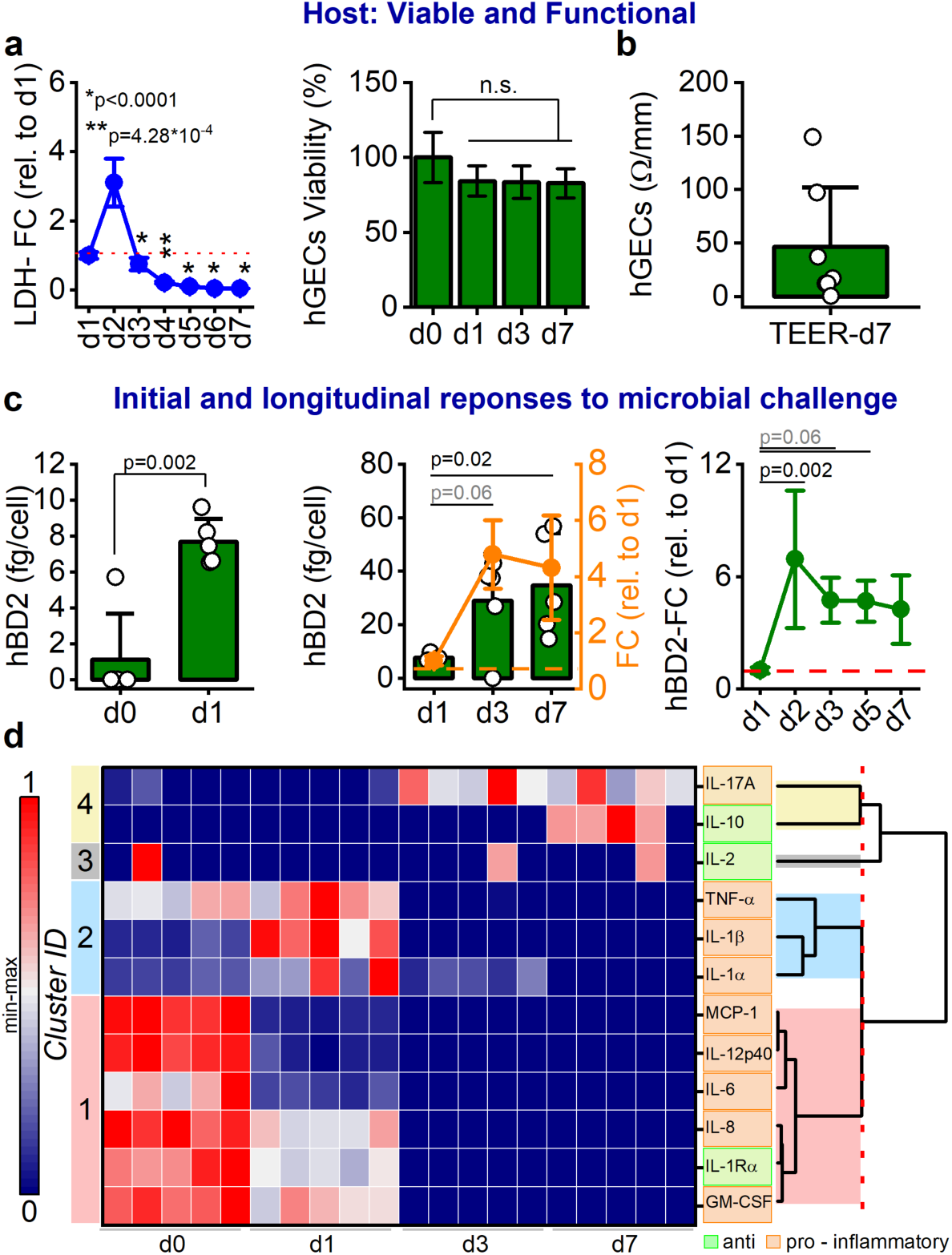
Dysbiotic microbiomes do not impair OTM viability and ability to respond to bacterial challenges. **a.** OTM viability and functional epithelial barrier response: Left — changes in LDH release relative to Day 1 (dashed red line) after subgingival microbiome inoculation. One-Way ANOVA repetitive measures with Dunnet (d1) post-hoc, p-values (<0.05) (n=5); the line plot represents the mean ± SD. Middle — DNA quantification via picoGreen analysis from gingival epithelial cells. The bar plot represents the mean ± SD (n=4), n.s. = not significant. Right — TEER assessments (Ω/mm) of oral epithelium on Day 7 (n=6). **b.** Initial and longitudinal responses of OTM to microbiome inoculation: Left — initial response (Day 0 – Day 1), hBD2 (fg/cell) release relative to oral epithelium cell density (n=5); Two-Sample Student’s t-test, p-value (<0.05). Middle — prolonged response (Day 1- Day 3 – Day 7), hBD2 release relative to oral epithelium cell density and fold change (orange dashed line) relative to Day 1 (n=5); One-Way ANOVA repetitive measures with Dunnet (d1) post-hoc, p-values (<0.05); the line graph represents the mean±SD. Right — longitudinal response (Day 2 – Day 7) fold change of ELISA data (fg) relative to day 1 (red dotted line) (n=5); One-Way ANOVA repetitive measures with Dunnet (d1) post-hoc, p-value < 0.05, the line graph represents the mean ± SD. **d.** Heat map showing the longitudinal inflammatory response of the OTM (pro-inflammatory and anti-inflammatory cytokines). Each cytokine (pg/cell) was rescaled with min-max normalization. Hierarchical clustering (right) identifies cytokine clusters, with a red dashed line indicating the cut-off value. Each cluster is color-coded as indicated (n=5).

### Preservation of microbial diversity and dysbiotic signature within the OTM

To evaluate the OTM’s capacity to support microbial colonization, growth, and diversity, subgingival plaque samples were inoculated into the OTM and taxonomic profiles were assessed at Day 0 (*h*plaque0, red) and Day 7 (native day 7, green) (**Figure 3a**). Relative abundance analysis confirmed the dysbiotic microbial signature in gingivitis-derived inocula (**Figure 3b**), with key genera, including *Campylobacter*, *Capnocytophaga*, *Fusobacterium*, *Prevotella*, *Selenomonas*, *Treponema*, and *Veillonella*, remaining prevalent at Day 7, supporting the long-term preservation of disease-associated microbial community [70]. OTU analysis revealed 175 shared taxa between baseline and Day 7 samples, with 90 and 9 OTUs unique to *h*plaque0 and native day 7, respectively (**Figure 3c**). Diversity indices indicated reduced richness and evenness over time, alongside community segregation relative to the initial inoculum (**Figure 3d-e**). Cladogram analysis (**Figure 3f**) identified enrichment of *Bacteroides*, *Actinobacteria*, and *Fusobacteria* in *h*plaque0 (red), whereas *Absconditabacteria* and *Proteobacteria* were enriched in OTM on Day 7 (green). To further resolve community structure, modularity analysis based on pairwise Spearman rank correlations (**Figure 3g**) identified three co-occurring modules, consistent with the shared and unique OTU distributions (**Figure 3c**). Module 1 exhibited the strongest and Module 3 the weakest intra-module correlation strength (**Supplementary Information**). The modularity value (0.206), used as a proxy for community compartmentalization, indicated a weak but detectable modular structure. This suggests a modest tendency for species to cluster into distinct co-occurring groups while maintaining moderate inter-module connectivity. Module 1 contained species present in both groups or uniquely present in native Day 7. Module 2 comprised species found in both groups or only in *h*plaque0. Module 3 included species shared across both groups.

**Figure 3:**
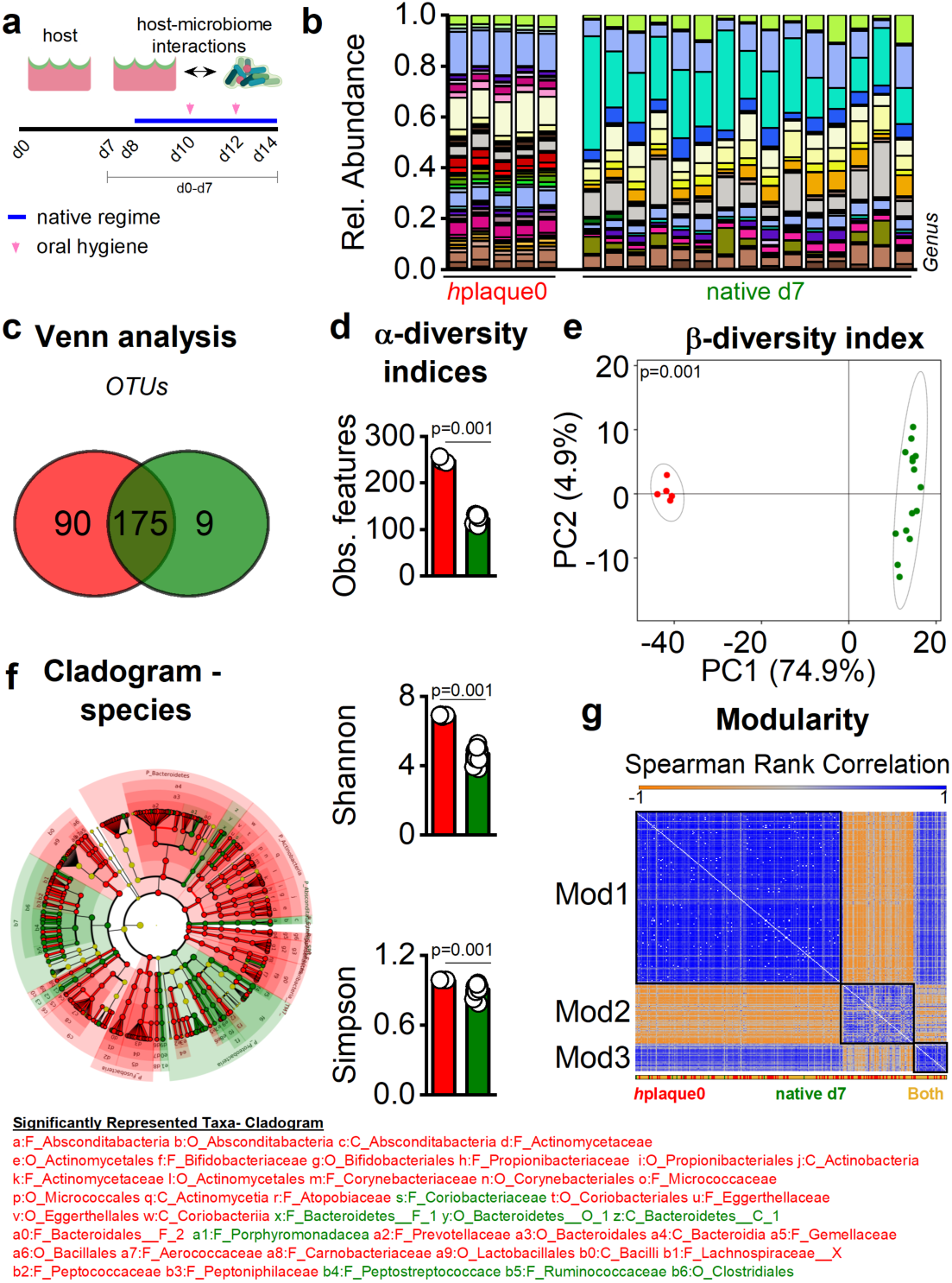
OTM promotes the colonization of dysbiotic microbiomes, supporting their growth and preserving most of the diversity. **a.** Schematic of host-microbiome interactions in the OTM, highlighting native regime (Day 1 to Day 7) and oral hygiene (Day 3 and Day 5) timeline. **b-g.** Comparison between hplaque0 (red – pooled subgingival plaque microbiomes, N-human subjects=20) and microbiomes cultured in the OTM (green, native d7, n=5): **b.** Relative abundance — Genus; legend is reported in **Supplementary** Figure 4**. c.** Venn analysis — species. **d-e:** α- and β-diversity indices — α: Observed features, Shannon and Simpson, Kruskal–Wallis H test, p-values (<0.05) are shown in the figure; single data points (circles) are shown in the graph; β: Principal component analysis (PCA) by means of Euclidean distance (Aitchison distance), PERMANOVA test, p-value (<0.05) is shown in the figure. **f.** Cladogram - Analysis of the differential abundance of statistically significant taxonomic levels (species). Color-code: red and green nodes indicate taxa with statistically significant differences in abundance between groups, while yellow nodes indicate non-significant taxa. The diameter of the circle is proportional to the relative abundance of each taxon. **g.** Heatmap representing similarity of the taxa (species) between hplaque0 and native d7 via Spearman Rank Correlation (−1;1). Black squares indicate a distinct module (**Supporting Information**). The color-coded horizontal bar indicates, for each species, whether the species was present exclusively in the hplaque0 (red), native d7 (green), or preserved over time (both conditions-yellow). Cartoon was made in Biorender.com.

### Gingival sulcus depth: spatial organization of dysbiotic microbial communities within the OTM

Dental plaque accumulation causes local changes at the gingival sulcus, such as inflammation or reduced tissue oxygenation [71–73] in gingivitis. Microbiome inoculation in the OTM supported statistical change (Day 0 – Day 1) in sulcus oxygenation, indicative of microbiome attachment and distribution (**Supplementary Figure 1c**), with differences in oxygen content among different regions of the OTM (Day 1: upper *vs*. middle region, p=0.03; upper *vs*. lower region, p=0.002; middle *vs.* lower region, p>0.05). Viability assessments by CLSM, flow cytometry, and SEM micrographs (**Supplementary Figure 2b-c**) on Day 7 confirmed the microbiome’s viability (85.3% live population *vs.* 13.9% dead population) and its distribution within the depth of the sulcus. To assess the spatial distribution of the microbiome within the OTM in relation to oxygen gradients [35], we performed taxonomic analyses for each region and compared differences and similarities (**Figure 4**). Each region shared approximately 55% of the OTUs with *h*plaque0 (**Supplementary Figure 3**). The relative abundance of the aerobic genera, including *Neisseria* [74] and *Stenotrophomonas* [75], was higher in the upper region (%: 13; 30) than in the middle (%: 10; 21) and lower (%: 5; 17) regions (**Figure 4a**). Conversely, anaerobic genera such as *Dialister* [76] and *Veillonella* [77] showed increasing abundance from the upper (%:0.8; 2.8) to the middle (%: 2.4; 4) and lower (%: 3; 9) regions. While approximately 73% of species (**Figure 4b**) were shared across all regions, indicating bacterial colonization, richness and diversity throughout the entire depth of the sulcus (**Supplementary Figure 2c** and **Figure 4c**), several taxa were region-specific (UR: 8.2%; MR: 4.3%; LR: 6.5%), with the upper and lower regions sharing less than 1% of total OTUs (**Figure 4d**). Differential abundance analysis (**Figure 4e-f**) identified region-enriched phylotypes (cladogram - **Figure 4e**), with *Actinobacteria* enriched in the upper region (green) and *Firmicutes* and *Proteobacteria* in the lower region (red). LEfSe (**Figure 4f**) further identified 20 and 56 statistically significant taxa (LDA score > 2) in the upper and lower regions, respectively. For example, a facultative anaerobic genus, *Aggregatibacter* [78], showed a decreasing gradient from upper to lower (6–3%) regions, whereas a facultative anaerobic genus, *Streptococcus* [79], was highly enriched in the lower region (16%) compared to the upper region (6%).

**Figure 4:**
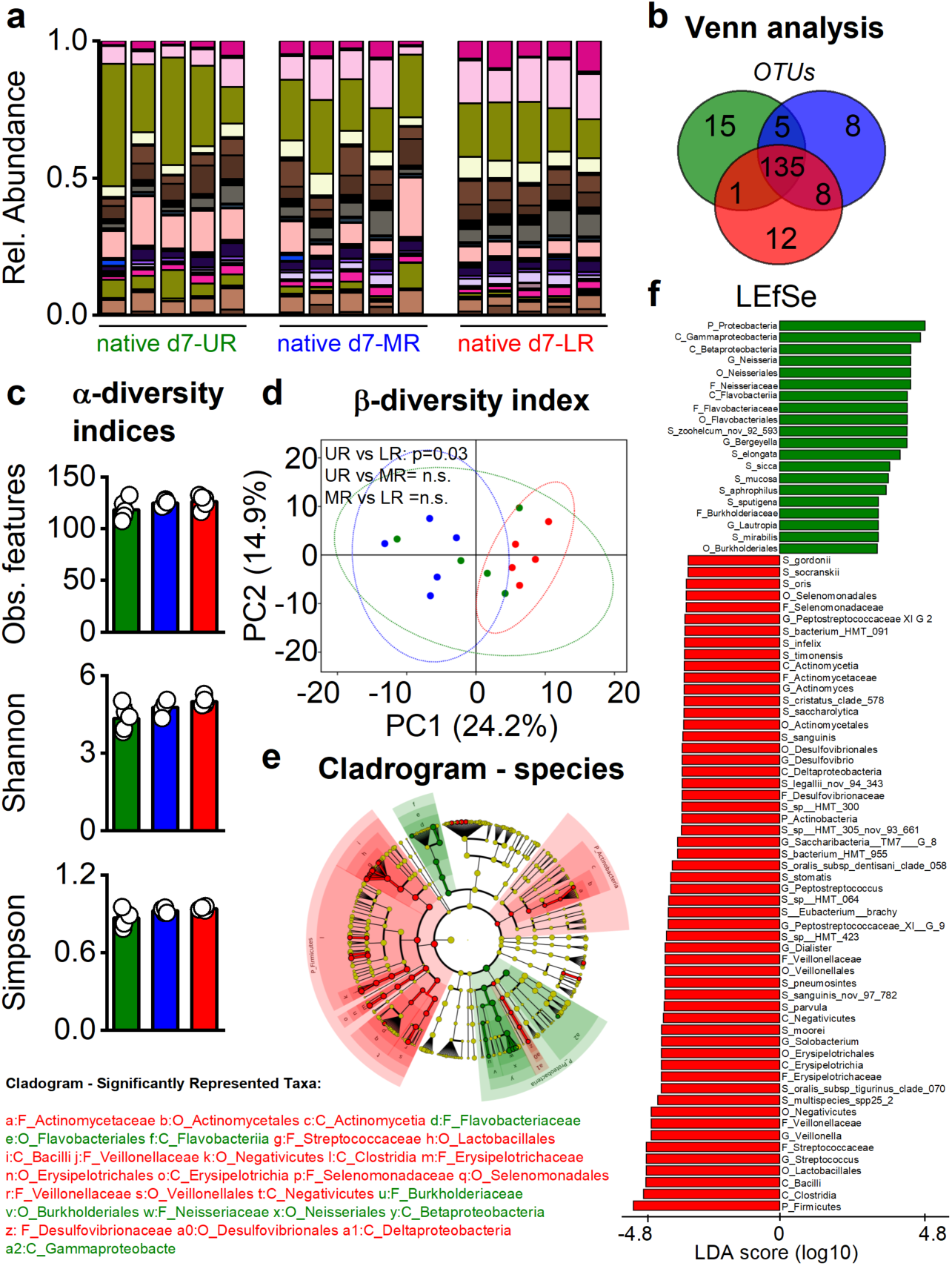
OTM favors organization of microbial communities according to their oxygen tolerance. **a-f.** Comparisons between gingival sulcus depths (regionality) in the OTM on Day 7: upper region (UR — green), medium region (MR — blue), lower region (LR — red), n=5: **a.** Relative abundance — Genus; legend is reported in **Supplementary** Figure 5**. b.** Venn analysis — species. **c-d:** α- and β-diversity indices — α: Observed features, Shannon and Simpson; single data points (circles) are shown in the graph; β: PCA by means of Euclidean distance (Aitchison distance), PERMANOVA test, p-value (<0.05); n.s.= not significant. **e.** Cladogram — Analysis of the differential abundance of statistically significant taxonomic levels (species). Color-code: red and green nodes indicate taxa with statistically significant differences in abundance between the UR and LR, while yellow nodes indicate non-significant taxa. The diameter of the circle is proportional to the relative abundance of each taxon. **f.** Linear discriminant analysis of the size effect (LEfSE) of statistically significant enriched taxa between the UR and LR.

### Gingival Crevicular Fluid (GCF) analysis in periodontal states

To evaluate the host’s inflammatory status at the time of microbiome sampling, GCFs (epithelium exudates) [80,81] were collected from periodontally healthy or gingivitis human subjects and subsequently analyzed. Biomarker profiling identified both pro- and anti-inflammatory cytokines, aligned with clinical diagnosis of each group (**Figure 5a**). Given the interconnected nature of signaling networks [82] and the limitations of single-analyte analyses, we applied MANOVA to assess whether the combined cytokine profile could discriminate between the two clinical states (healthy *vs.* gingivitis). MANOVA revealed significant differences between healthy and gingivitis groups (Wilks’ Lambda=0.368, F(11,26)=4.05, p=0.002, partial η^2^=0.632). Linear discriminant analysis (LDA) further identified key contributors to group separation. The first canonical variate accounted for 79.5% of the explained variance and showed significant separation between the gingivitis and healthy groups (p=0.001; **Figure 5b**), suggesting distinct cytokine signatures. Posterior probability analysis confirmed classification performance, with most samples assigned to their respective clinical groups, although partial overlaps were observed (**Figure 5c**). Ranking cytokines by normalized weighted contribution to group discrimination (**Figure 5d**) identified IL-1α as the strongest discriminator, followed by IL-8 and IL-1β. MCP-1, IL-12p40, GM-CSF, and IL-17A contributed moderately, whereas IL-10, IL-2, TNF-α, and IL-6 had minimal influence. Collectively, these findings indicate that pro-inflammatory cytokines, particularly members of the IL-1 family, primarily drive discrimination between the two periodontal states and define a subset of biomarkers for oral tissue status identification.

**Figure 5:**
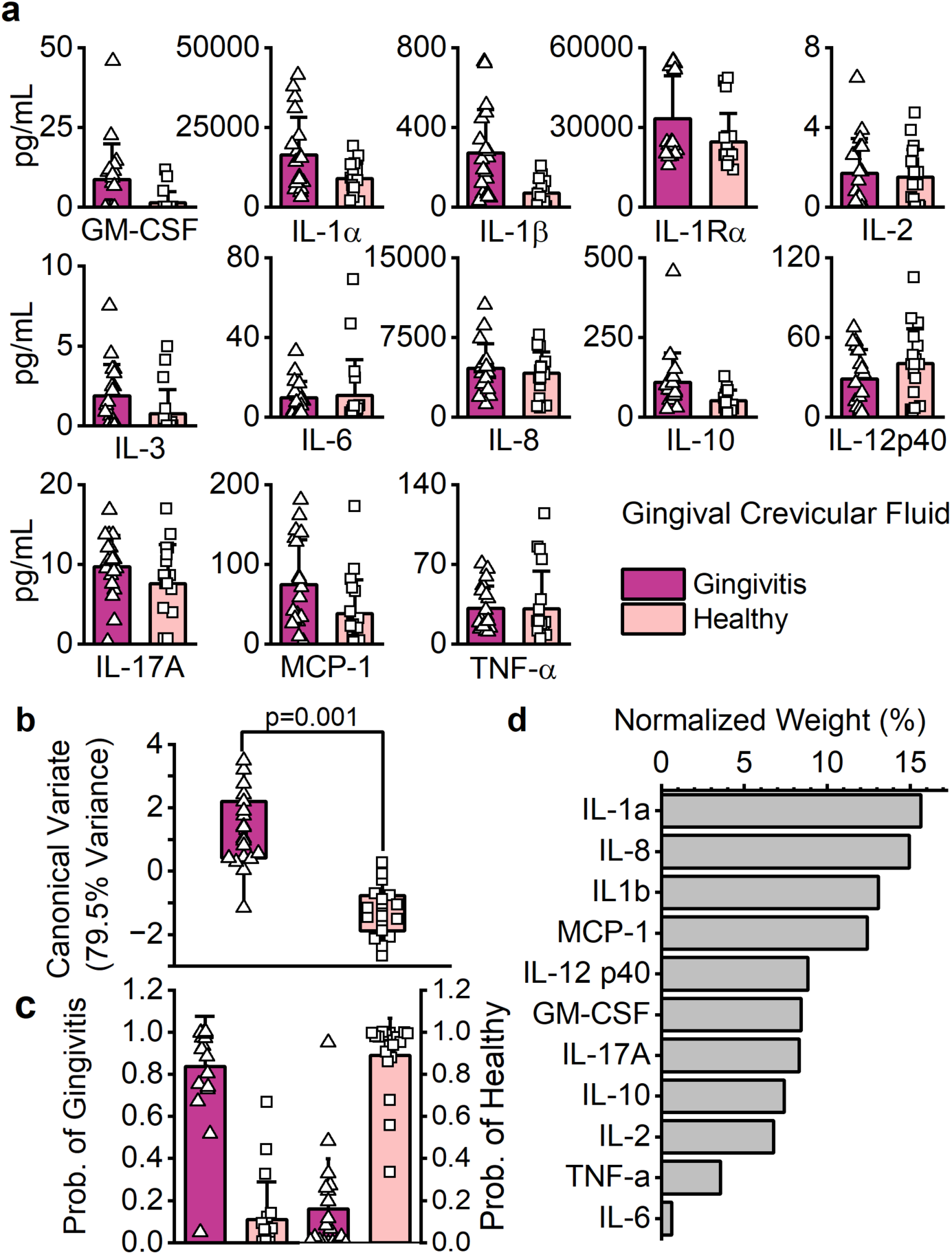
Gingival crevicular fluid (GCF): a report card of periodontal health. **a.** Pro- and anti-inflammatory cytokines (pg/mL) in GCF isolated from gingivitis (dark pink, triangles) and healthy (pink, squares) subjects (n=19 human subjects/clinical state) and quantified via Milliplex® assay; single data points (triangles and squares) are reported in the graphs. **b.** Box plots of the first canonical variate scores (79.5 % of total explained variance) derived from linear discriminant analysis (LDA); single data points (triangles and squares) are reported in the graph. **c.** Predicted probabilities for each patient (data point) of belonging to one of the two groups based on their individual cytokine profile. The left panel shows the probability of belonging to the gingivitis group, while the right panel shows the probability of belonging to the healthy group; single data points (triangles and squares) are reported in the graphs. **d.** Cytokines ranked (normalized weight (%)) in descending order of importance based on their weighted contribution to group discrimination in LDA.

### OTM: an advanced tissue engineering platform for periodontal clinical calibration

The OTM enables identification of early biomarkers of oral tissue inflammation while accounting for inter-donor variability [83] (**Figure 6**). A consistent methodology was used to extract and process epithelial exudates (GCF and OTM) (**Figures 1** and **6a**). The longitudinal profiles of IL-1α, IL-1β, IL-8, and MCP-1 in the OTM were analyzed based on cytokine ranking (**Figure 5d**) across clinical conditions (**Figure 6b**). Proinflammatory mediators IL-1α, IL-1β, and MCP-1 were upregulated in the gingivitis OTM (dark pink) compared with the healthy OTM (pink) on Day 1. Cytokine levels decreased by Days 3 and 7, indicating an early modulatory response followed by stabilization. No difference in IL-8 cytokine release was detected (p>0.05). The hBD2 profile (**Figure 6c**) in GCF showed higher concentrations in gingivitis compared to healthy subjects, consistent with inflammatory response to plaque accumulation [84]. In contrast, hBD2 secretion in the OTM was higher under healthy conditions versus inflamed (gingivitis) ones on Day 1, with a significant effect of time in gingivitis conditions.

**Figure 6:**
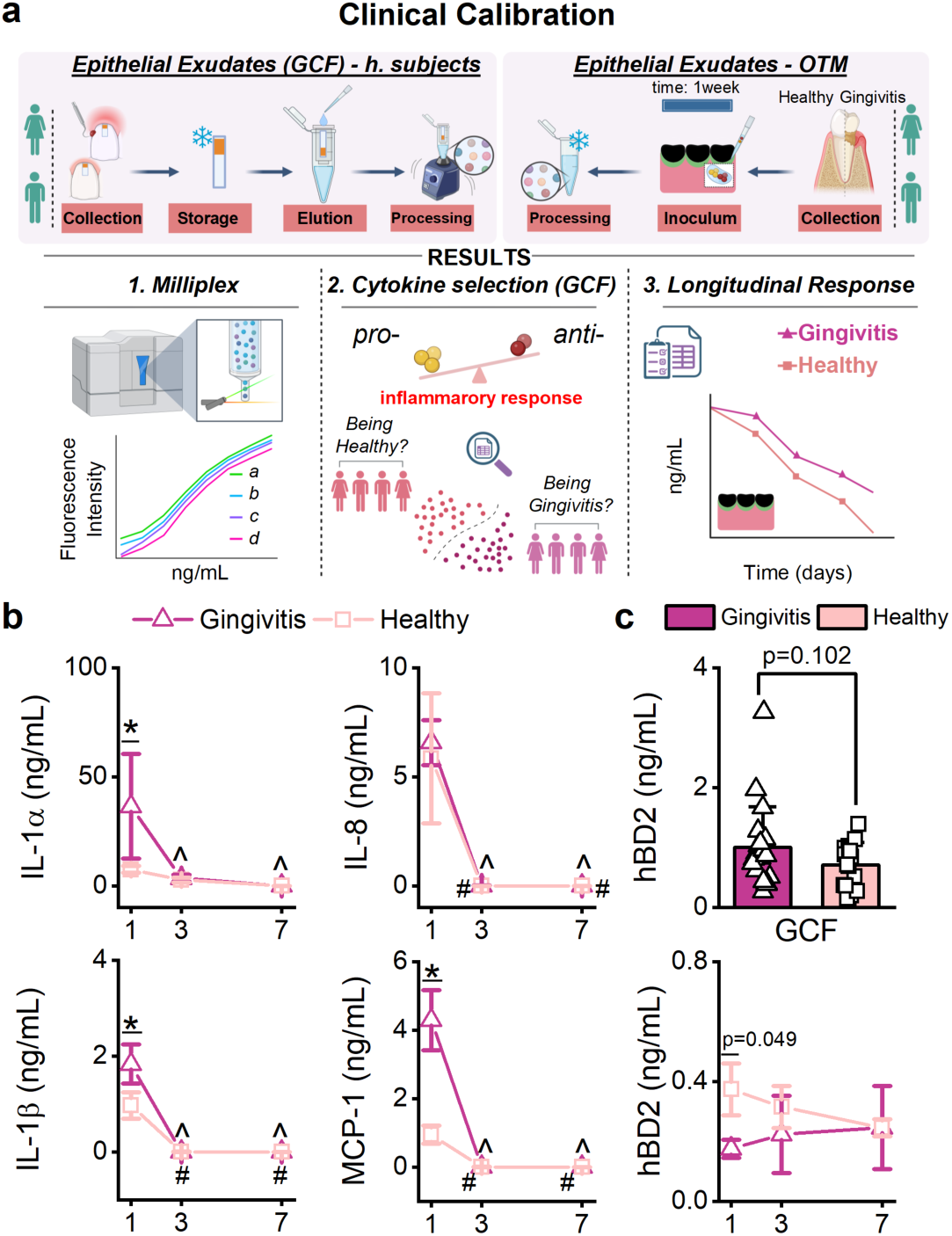
Clinical calibration indicates an early pro-inflammatory response OTM under gingivitis conditions. **a.** Clinical calibration workflow: epithelial exudates from GCF (n=19 human subjects/clinical state) and OTM (n=5) were isolated and processed using similar protocols, before being quantified using the Milliplex® assay. Subsequently, statistical analysis (Figure 5c) of the pro-inflammatory and anti-inflammatory response on GCF was used to select four cytokines (IL-1α, IL-1β, IL-8, and MCP-1) with the highest normalized weight (%) in discriminating host response based on clinical status. The longitudinal profile of these cytokines was then evaluated in the OTMs. **b-c.** OTM pro-inflammatory response based on inoculum of healthy (n=5, n=8 healthy subjects) [37] or gingivitis (n=5, n=20 gingivitis subjects) subgingival microbiomes. **b.** Line plots representing the mean ± SD expressed as ng/mL of IL-1α, IL-1β, IL-8, and MCP-1 in gingivitis (dark pink, triangles) and healthy (pink, squares) OTMs. Two-Way repetitive measures ANOVA, with Bonferroni post-hoc; p-values (<0.05) are reported in **Supplementary Table 2**; asterisks (*) represent statistical significance (p<0.05) between disease states for each time point; carrots (^ - gingivitis) and hashtags (# - healthy) represent statistical significance (p<0.05) within the same condition, overtime, compared to d1. **c.** hBD2 expression (ng/mL) in: top — GCF, between healthy and gingivitis subjects (n=19 human subjects/clinical status). Two-Sample Student’s t-test with Welch correction (unequal variance, p=0.01), p-value (<0.05) is shown in the figure; single data points (triangles and squares) are reported in the graph. Bottom — OTMs (n=5/clinical status); the line plot represents the mean ± SD expressed as ng/mL; Two-Way repetitive measures ANOVA, with Bonferroni post-hoc; p-values (<0.05). Cartoon was made in Biorender.com.

### Characterization of subgingival microbial communities based on clinical state

To assess the OTM’s ability to support microbiome persistence under different clinical states, we compared taxonomic profiles for each condition (**Figure 7**). Relative abundance assessments (**Figure 7a**) demonstrated that the OTM maintains subgingival plaque microbiomes from both healthy and gingivitis patients over 7 days. Gram-positive genera *Rothia* and *Gemella* were less abundant in the healthy inoculum than in the dysbiotic microbiome; however, the commensal *Gemella* [85] showed increased prevalence in healthy OTM by Day 7. *Actinomyces,* a commensal genus with opportunistic pathogenic potential [86][87], was more abundant in gingivitis than in healthy microbiomes (*h*plaque0), but was reduced (gingivitis) or undetected (healthy) by Day 7. Among Gram-negative genera, *Porphyromonas* and *Treponema* were more abundant at the time of inoculum in gingivitis conditions, with the gingivitis OTM supporting *Porphyromonas* enrichment (% - *h*plaque0: 2.45; native d7: 6.04). Lastly, the abundance of the *Tannerella* and *Fusobacterium* genera were similar in both clinical conditions on *h*plaque0; however, the relative abundance of *Fusobacterium* decreased more markedly in healthy OTMs (%: *h*plaque0: 4.9; native d7:1.6) than in gingivitis (%: d0: 5.9; d7:3.6) over 7 days. Venn analysis (**Figure 7b**) indicated that only 14.8% of the total OTUs were shared between the initial plaque samples from healthy and gingivitis donors; specifically, the inflamed state exhibited greater complexity compared to healthy conditions (**Figure 7c**). Consistent with previous findings [37], diversity and richness (α-diversity indices) were decreased from Day 0 (*h*plaque0) and Day 7 (native d7) in both clinical states, while most of the diversity was preserved (**Figure 7c**). Segregation from the original samples was also indicated by the β-diversity index (**Figure 7d**) based on clinical periodontal condition (*h*plaque0-native d7). In addition, greater segregation (**Figure 7d**) was observed among OTMs on Day 7 in both conditions (OTM*G*_day 7 *vs.* OTM*H*_day 7), with only 1.7% of species shared (**Figure 7b**).

**Figure 7:**
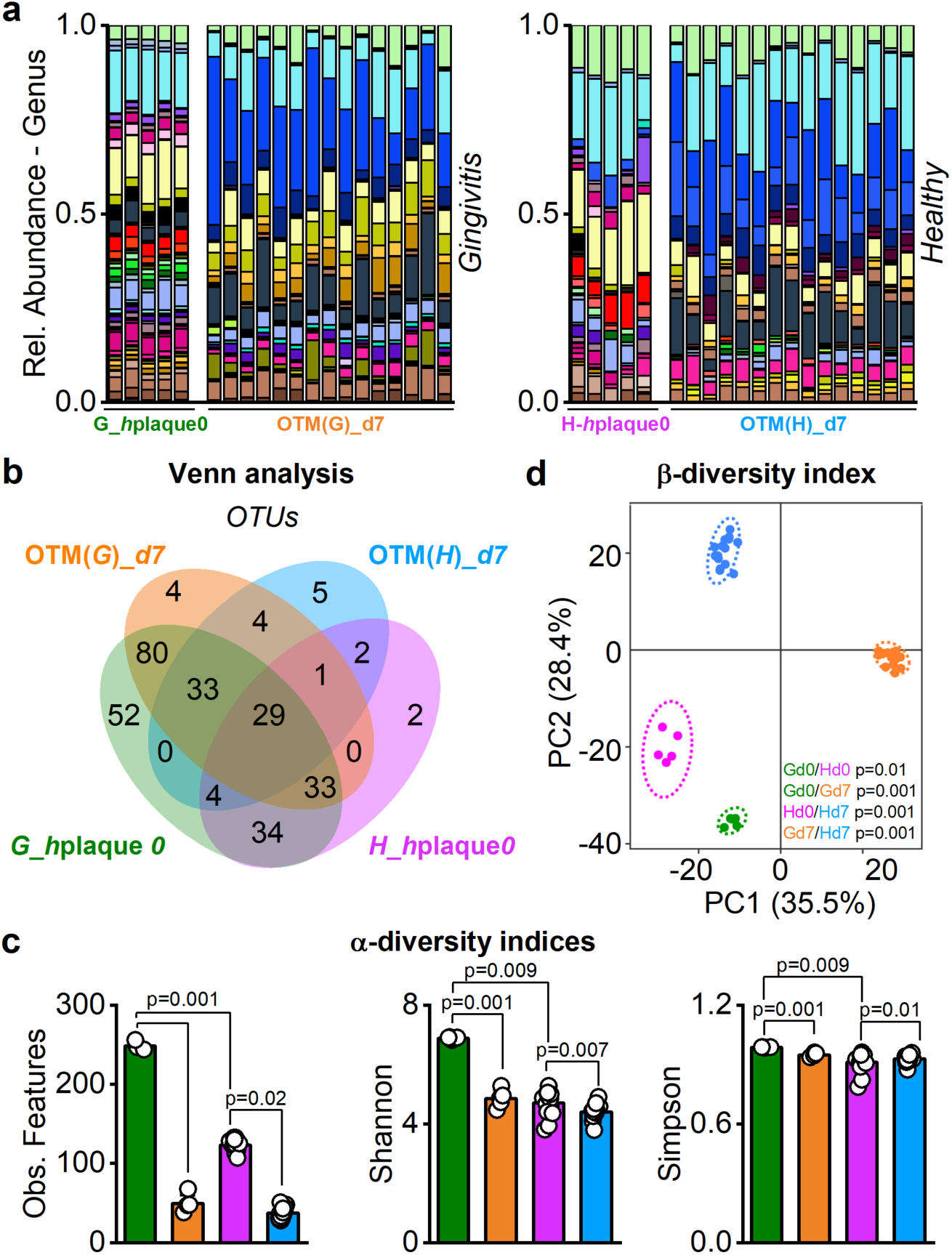
OTM can preserve disease state microbial signature long-term. **a-d.** Comparisons between hplaque0 (healthy, purple – N-human subjects=8), hplaque0 (gingivitis, green – N-human subjects=20), OTMd7 (healthy, blue, n=5) [37] and OTMd7 (gingivitis, orange, n=5): **a.** Relative abundance — Genus; legend is reported in **Supplementary** Figure 6**. b.** Venn analysis — species. **c-d:** α- and β-diversity indices — α: Observed features, Shannon and Simpson, Kruskal–Wallis H test, p-values (<0.05); β: Principal component analysis (PCA) by means of Euclidean distance (Aitchison distance), PERMANOVA test, p-value (<0.05) are shown in the figure.

### Ionic and Metabolic response in OTM based on microbial inocula

Leveraging the potential of saliva for early detection of periodontal inflammation [88], we assessed longitudinal changes in biomarkers for carbon metabolism (glucose, lactate, glutamine and glutamate) [65,66], extracellular ionic composition (sodium, potassium, and calcium) [66], and nitrogen metabolism (ammonium) [89] (**Figure 8**). In gingivitis, glutamine concentrations steadily decreased over time, whereas they remained consistent in healthy conditions, with no differences observed in glutamate profiles. Glucose was consumed under all conditions, consistent with cellular substrate utilization, while lactate levels declined over time. Sodium, potassium, and calcium exhibited similar trends, increasing under healthy conditions but decreasing in gingivitis. Nitrogen metabolism was enhanced in gingivitis, suggesting enhanced deamination and amino acid catabolism under dysbiotic conditions [90]. Finally, pH dynamics (**Figure 8b**) revealed initial acidification following microbiome inoculation (Day 0 through Day 1), followed by progressive alkalinization (Day 2-7) in both clinical conditions.

**Figure 8:**
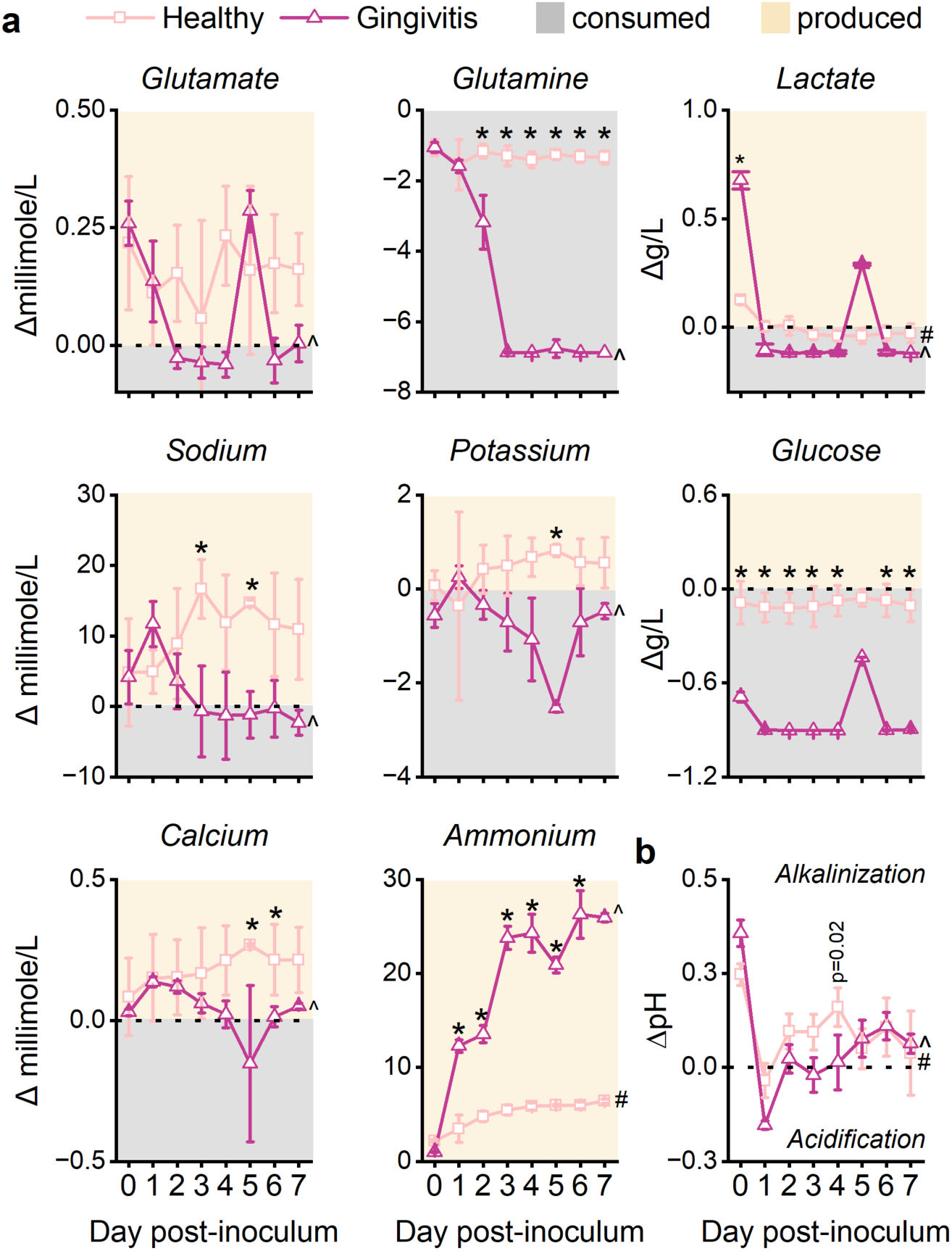
OTM modulates ionic and metabolic response based on microbial signature. **a.** Measurements of salivary metabolic indexes in healthy (n=5) and gingivitis (n=5) conditions over a 7-day period. Line graphs represent the mean ± SD, with shaded regions separated by a black dashed line (0) denoting net production (yellow) or consumption (grey) relative to the artificial saliva baseline (background). Two-Way repeated-measures ANOVA with Bonferroni post-hoc; single p-values (p<0.05) are reported in **Supplementary Table 3;** asterisks (*) represent the statistical significance between groups at the same time point; One-way repeated measures ANOVA with Dunnett (d0) post-hoc test was performed to assess statistical differences (p<0.05) within the same condition (carrots (^), gingivitis; hashtags (#), healthy) relative to d0; single p-values (p<0.05) are reported in **Supplementary Table 3**. **b.** Longitudinal pH measurements relative to artificial saliva pH (7.4, black dashed line) in OTMs exposed to healthy [37] (n=3) or gingivitis (n=5) subgingival plaque microbiomes. Two-Way repeated-measures ANOVA with Bonferroni post-hoc was performed to identify statistical differences (p<0.05) between group at the same time point is reported in the figure; One-Way repeated measures ANOVA with Dunnett (d0) post-hoc test was performed to assess statistical differences (p<0.05) within the same condition (carrots (^), gingivitis; hashtags (#), healthy) relative to Day 0; single p-values are reported in **Supplementary Table 4**.

## Discussion

The mechanisms underlying the initiation and progression of periodontal inflammatory diseases, including gingivitis and periodontitis, remain poorly defined. This knowledge gap reflects the complex and pronounced interindividual variability of microbial community dynamics, host immune responses, and host-microbiome crosstalk, which together hinder the identification of early disease drivers [13]. To overcome these challenges, physiologically relevant *in vitro* models offer a controlled platform to dissect early tissue responses to dysbiotic microbiomes [14,27].

In a previous study, we established that sustained host–microbiome interactions under periodontal health conditions in an *in vitro* oral tissue model (OTM) require integration of key physiological parameters, including cyto-anatomical architecture, native shear stress, saliva buffering, and tightly regulated oxygen and pH gradients [35,37]. Building on this foundation, the present study exposed mature anatomical models to dysbiotic microbiomes derived from patients with gingivitis and longitudinally evaluated host and microbial responses, including cell viability, pro- and anti-inflammatory signaling, and microbial community composition. In addition, model-derived inflammatory, metabolic, and microbial readouts were benchmarked against patients-derived GCF and subgingival microbiomes to enable clinical validation [91,81,92].

Gingivitis is initiated by plaque accumulation at the tooth-gingiva interface [93] accompanied by a shift from a eubiotic to a dysbiotic microbiome that triggers localized inflammation and host immune activation [2,94,95]. Accordingly, we inoculated a mature anatomical gingival model with human subgingival plaque microbiomes isolated from patients with gingivitis to investigate early host–microbiome interactions. The use of dysbiotic polymicrobial communities for *in vitro* inflammatory stimulation is consistent with the polymicrobial synergy and dysbiosis (PsD) hypothesis, which posits that periodontal conditions arises by reciprocally reinforcing interactions between dysregulated host inflammatory responses and dysbiotic microbiomes [8]. In contrast, most existing *in vitro* models induce inflammation using lipopolysaccharide (LPS) [96,97], pro-inflammatory cytokine cocktails (such as IL-1β and TNF-α) [32,33], or in combination, typically eliciting acute inflammatory responses over short time frames (24–48 h). While these reductionist approaches have provided valuable insights into discrete signaling pathways, they fail to recapitulate the complexity of inflammatory and metabolic signaling, as well as the dynamic bacteria-bacteria and bacteria-host interactions that characterize *in vivo* periodontal diseases [8,13]. The gingival tissue is continuously exposed to microbial stimuli, requiring both barrier function and rapid immune responsiveness [82,98]. Accordingly, we first assessed the long-term OTM viability and barrier integrity under dysbiotic challenge, considering its exposure to microbial metabolic byproducts [27]. The model exhibited sustained viability and preserved functional epithelial structure (**Figure 2a-b** and **Supplementary Figure 1a-b**). Following inoculation, the OTM mounted a dynamic antimicrobial and inflammatory response. The antimicrobial peptide hBD2, a key epithelial defense factor targeting both Gram-positive and Gram-negative bacteria [99,100], was robustly induced and remained elevated throughout the culture period (**Figure 2c**). This response is consistent with known inducers such as TNF-α [100], and *Fusobacterium nucleatum* [103,99], both of which were present and elevated at early time points (Days 0-1) (**Figure 2** and **3**). Cytokine profiling further revealed temporally structured responses, with distinct modules identified by hierarchical clustering analysis: Module 1 (MCP-1, IL-12p40, IL-6, IL-8, IL-1Rα, GM-CSF), Module 2 (TNF-α, IL-1α, and IL-1β), Module 3 (IL-2), and Module 4 (IL-10, IL-17A) (**Figure 2d**). Module 1 consisted of pro-inflammatory cytokines, except for IL-1Rα (anti-inflammatory). These cytokines were mostly upregulated on Day 0 and slightly decreased (IL-8, IL-1Rα, GM-CSF) or not expressed (MCP-1, IL-12p40, IL-6) on Day 1. Opposite trend (no expression on Day 0, upregulation on Day 1) was instead observed for the pro-inflammatory cytokines in Module 2 (TNF-α, IL-1α, and IL-1β). As a primary line of defense upon microbial challenge, pro-inflammatory mediators of IL-1 (α-β) and TNF families are secreted by gingival (epithelial and fibroblast) cells [82]. Consistent with these observations, pro-inflammatory cytokines were upregulated in the OTM on Day 1. Increased expression of TNF-α and IL-1β by gingival keratinocytes has been previously associated with hBD-2 secretion [104], which was likewise observed at Day 1. Interestingly, although IL-1α levels declined by Day 3, this cytokine has been implicated in promoting IL-17A production (Module 4), a response that was sustained and significantly upregulated in the OTM from Day 3 through Day 7. IL-17A is pivotal in immune surveillance and in preserving tissue homeostasis at mucosal interfaces [105]. However, while serving a protective role, prolonged release of IL-17A has been associated with severe gingival inflammation, tissue damage, and bone loss [106]. Consequently, release of anti-inflammatory IL-2 (Module 3) and IL-10 (Module 4) was found to be upregulated in the OTM on Day 3 to Day 7, suggesting a compensatory regulatory response rather than activation of a pro-inflammatory pathway [82]. Collectively, these findings demonstrated that OTM withstands dysbiotic microbial challenges while preserving viability, barrier integrity, and dynamically regulating inflammatory responses, supporting its utility as a platform to interrogate early mechanisms of periodontal disease.

In our model, microbiome inoculation (Day 0 to Day 1) resulted in reduced tissue oxygenation (**Supplementary Figure 1c**) with values (O_2_>2%) comparable to clinical measurements at periodontal pockets [71]. By Day 7, the OTM supported high microbial viability (85.3% live microbial population) (**Supplementary Figure 3**) and preserved overall community diversity, despite shifts in diversity indices (**Figure 3**). Clinically relevant taxa—including *Streptococcus oralis*, *F. nucleatum,* and *Tannerella forsythia*—were detected, consistent with their role as early colonizers (*S. oralis*), bridging species (*F. nucleatum*), and inflammation-associated pathobionts (*F. nucleatum and T. forsythia*) [107–109,8,110]. Modularity analysis further resolved community organization, identifying *S. oralis* in Module 1 and *F. nucleatum* and *T. forsythia* in Module 2 at Day 7 (**Figure 3** and **Supplementary Information**), supporting their established role in early biofilm development and polymicrobial synergy. The OTM also preserved interspecies interactions, including the association between *Veillonella parvula* and *Porphyromonas gingivalis*, which were co-grouped in Module 1. Consistent with reports that *V. parvula* promotes the overgrowth of low-density *P. gingivalis* [111], we observed concurrent increases in both species from the initial inoculum (*h*plaque0, Day 0) to Day 7 (*V. parvula*: 1.4% to 1.9%; *P. gingivalis*: 0.2% to 5.35%). These coordinated shifts support the persistence of interspecies interactions within the OTM and underscore its capacity to maintain dysbiotic community dynamics over time (**Figure 3** and **Supplementary Information**). Lastly, consistent with our previous findings [35,37], we observed a spatial stratification of the microbiome along the pocket depth, with the upper region differing significantly from the lower region of the OTM (**Figure 4** and **Supplementary Figure 3**). Collectively, these findings demonstrate that the OTM supports colonization and growth of subgingival plaque communities throughout the full pocket depth, driven by the establishment of physiologically relevant oxygen gradients, while preserving the diversity and taxonomic composition characteristic of periodontal disease.

From an inflammatory perspective, defining the onset of gingivitis remains challenging due to weak correlations between clinical manifestations and molecular signature, compounded by high interindividual variability [112,113]. Gingivitis clinical diagnosis relies on probing, *e.g.* BoP index ≤30% and PI >1 with no attachment loss or tissue damage [20,47–49]. However, these diagnostic thresholds are operator-dependent and do not fully capture disease variability, as some patients exhibit extensive BoP despite minimal plaque deposits [114]. This suggests that dental plaque alone does not fully explain the disease etiology. Moreover, similar plaque levels can elicit different inflammation responses, with subjects classified as high responders (pronounced inflammation) or low responders (minimal inflammation) [17,113,114]. One of the main advantages of using *in vitro* models is the ability to have a controlled environment, enabling precise modulation of individual variables and facilitating the identification of specific inflammatory markers [14]. To initially correlate the composition of the subgingival plaque microbiomes with the degree of inflammation at the time of inoculation in the OTM between periodontally healthy and diseased subjects, we investigated inflammation markers in GCF and performed a 16S rDNA analysis (**Figures 5-7**). In agreement with clinical reports [115,113,17,116], cytokine profiles exhibited substantial variability across disease states (**Figure 5a**). Given the interdependency between pro- and anti-inflammatory cytokines [82], we employed a multivariate analysis of variance (MANOVA) to obtain an integrated assessment of cytokine response patterns. Our analysis indicated IL-1 α/β, IL-8 and MCP-1 cytokines can be adopted as predictors for disease onset and progression (**Figure 5d**). Comparisons between OTMs inoculated with eubiotic and dysbiotic microbiomes indicated increased concentrations of IL-1α/β, MCP1 and hBD2, but not IL-8, only on Day 1 upon dysbiotic microbiome exposure (**Figure 6**). This is in agreement with clinical evaluations in GCF, where higher amount of IL-1β has been detected in patients with gingivitis and periodontitis compared to healthy subjects [117,118,113]. However, reported IL-1β profiles are not consistently concordant between gingivitis and periodontitis cohorts [119–121], with concentrations decreasing or returning to baseline depending on oral hygiene treatments [121]. Similar variability has also been reported for IL-8, MCP1 and hBD2 across studies [84,113,115,122,123]. Although the OTM recapitulated clinical inflammatory patterns at early time points (Day 1), no significant differences were observed at later stages (Day 3 to Day 7), during which the host response shifted toward an anti-inflammatory phenotype rather than a sustained pro-inflammatory state, irrespective of the inoculum type (**Figure 2**) [37]. This temporal pattern suggests that prolonged microbial exposure may be required to fully reproduce gingival inflammatory trajectories in the OTM, consistent with clinical experimental gingivitis models that typically span 2-4 weeks [17,94]. Overall, inflammatory profiling reveals that the OTM mounts a time-dependent, inoculum-sensitive response, characterized by early pro-inflammatory activation followed by regulatory modulation.

Alongside inflammatory profiling, we assessed microbial composition in healthy and inflamed OTMs (**Figure 7**). Clinically, biofilm maturation in gingivitis conditions is accompanied by a shift in microbial composition [70]. In agreement with previous reports [124,125], gingivitis inocula displayed greater complexity than healthy samples. While both OTMs exhibited their expected profiles (healthy and dysbiotic), inflamed OTMs displayed greater heterogeneity, reflected by reduced similarity in β-diversity compared to healthy counterparts (**Figure 7d**). This divergence indicates differential responses of the model to microbiomes associated with distinct periodontal states. Notably, while pro-inflammatory responses peaked at Day 1, β-diversity progressively increased by Day 7 under dysbiotic conditions suggesting sustained remodeling of microbial community structure over time. Similar patterns have been reported in cross-sectional observational studies, where increasing microbial dissimilarity (greater distance) correlates with periodontal disease severity (gingivitis and periodontitis (I-II and III-IV)) [70]. Collectively, these findings support the ability of the OTM to preserve microbial diversity and maintain disease-associated microbial signatures characteristic of distinct clinical states.

Lastly, although microbial profiling data were not collected prior to Day 7, we profiled ionic and metabolic responses across periodontal states (**Figure 8**), which are widely used to assess microbiome behavior and host responses during periodontitis progression [126–129,88,130]. Metabolite analysis revealed temporal shifts aligned with clinical microbial signature, with negative (decrease) values indicating biomarker consumption and positive (increase) values indicating biomarker production. Inflamed OTM exhibited accelerated depletion of key carbon sources (glucose, glutamine, glutamate) alongside increased nitrogen substrates. This shift is consistent with enhanced amino acid catabolism and ammonium production, which may contribute to environmental alkalization in both conditions [131]. Additionally, lactate levels declined over time in both conditions, suggesting metabolic cross-feeding within microbial communities [132], consistent with taxa such as *Veillonella* that utilize lactate to support cooperative metabolism [13,133]. Sodium, potassium, and calcium displayed similar trends, increasing in healthy conditions but decreasing in gingivitis state, in line with clinical observation [134]. Reduced ion levels in the inflamed OTM likely reflect the absence of dentition, given that elevated ion concentrations in GCF and saliva are associated with alveolar bone degradation and impaired ion reabsorption during severe inflammation [127,134]. Despite reports that salivary ions can attenuate β-defensin activity by interfering with peptide-bacteria interactions [99], hBD2 levels remained stable or elevated from Day 1 to Day 7, indicating preserved host antimicrobial responses in healthy OTM. Overall, these metabolic and ionic profiles demonstrate the OTM’s capacity to distinguish periodontal states and capture the dynamic interplay between microbial activity and host responses.

Nevertheless, several limitations of the OTM should be acknowledged. The current model does not recapitulate the full structural complexity of the periodontium and does not incorporate neutrophils, which play a central role in gingivitis onset [95]. Moreover, although emerging evidence highlights sex dimorphism in periodontal conditions [135–138], gender was not considered as a variable in the present work, with healthy and dysbiotic subgingival microbiomes pooled across donors. While the OTM exhibited distinct responses to healthy and dysbiotic microbiomes at early time points (Day 1), responses converged toward an anti-inflammatory profile at later stages (Day 3-7).

## Conclusions

In this study, we further validate our previously developed oral tissue model (OTM) [35–37] as a physiologically relevant platform capable of sustaining both periodontal health–associated and dysbiotic microbiomes for up to seven days. The model preserves host viability, supports dynamically regulated inflammatory responses, and maintains microbial diversity and phenotype over time. Notably, this work establishes a clinical calibration against gingival crevicular fluid (GCF), for the first time in an *in vitro* system, where OTM responses capture inoculum-dependent inflammatory signatures, increased microbial dissimilarity under dysbiotic conditions, and coordinated host–microbiome metabolic interactions. While extending culture duration to better align with experimental gingivitis models and incorporating advanced molecular approaches (e.g., FISH and RNA gene sequencing) will further refine mechanistic resolution [94,139], our findings demonstrate that the OTM recapitulates key features of periodontal health and inflammation. As such, this platform provides a robust and translationally relevant system for interrogating early disease mechanisms and informing the development of predictive diagnostics and preventive therapeutic strategies.

## Supporting information

Supplementary Information_Community ID

## List of Abbreviations

OTM: Oral Tissue Model
D.I. water: Deionized Water
hGFCs: human Gingival fibroblast cells
hGECs: human Gingival Epithelial Cells
ALI: Air Liquid Interface
LDH: lactate dehydrogenase
TEER: TransEpithelial Electrical Resistance
gDNA: genomic DNA
GCF: Gingival Crevicular Fluid
PBS: Phosphate Buffer Saline
GM-CSF: Granulocyte-macrophage colony-stimulating factor
IL-1α: Interleukin 1 alpha
IL-1β: Interleukin 1 beta IL-1RA Interleukin 1 Receptor Antagonist
IL-2: Interleukin 2
IL-6: Interleukin 6
IL-8: Interleukin 8
IL-10: Interleukin 10
IL-12p40: Interleukin 12p40
IL-17A: Interleukin 17A
MCP-1: Monocyte Chemoattractant Protein 1
TNF-α: tumor necrosis factor-alpha.

## Declarations

### Ethics approval and consent to participate

Studies involved human subjects were conducted in compliance with the approved protocols: FIRB# 18-06, ADA Forsyth Institutional Review Board and IRB—20-090, the University of Massachusetts Lowell. Prior to specimen collections, all subjects signed the FIRB-approved informed consent.

### Consent for publication

Not applicable.

### Availability of data and material

The main data generated and analyzed during this study are included in this published article (and its supplementary information files). Any requests for raw data generated in the present manuscript should be directed at the corresponding author.

### Authors’ contributions

Conceptualization, project administration and funding acquisition: C.E.G. Investigation, methodology, validation and formal analysis: M.A., M.B., G.E.C., A.R.D., C.E.G; Data curation and visualization: M.A., M.B. G.E.C., A.R.D., C.E.G; writing — original draft preparation: M.A., M.B. G.E.C., A.R.D., C.E.G; writing — review and editing: M.A., M.B. G.E.C., A.R.D., C.E.G, H.H., B.J.P., X.H. All authors have read and agreed to the published version of the manuscript.

## Acknowledgements

We would like to thank Tsute Chen, Ph.D., for his help in navigating the sequencing reports. We are also thankful to all the students who have contributed to this project.

## Supplementary Figures

**Supplementary Figure 1:**
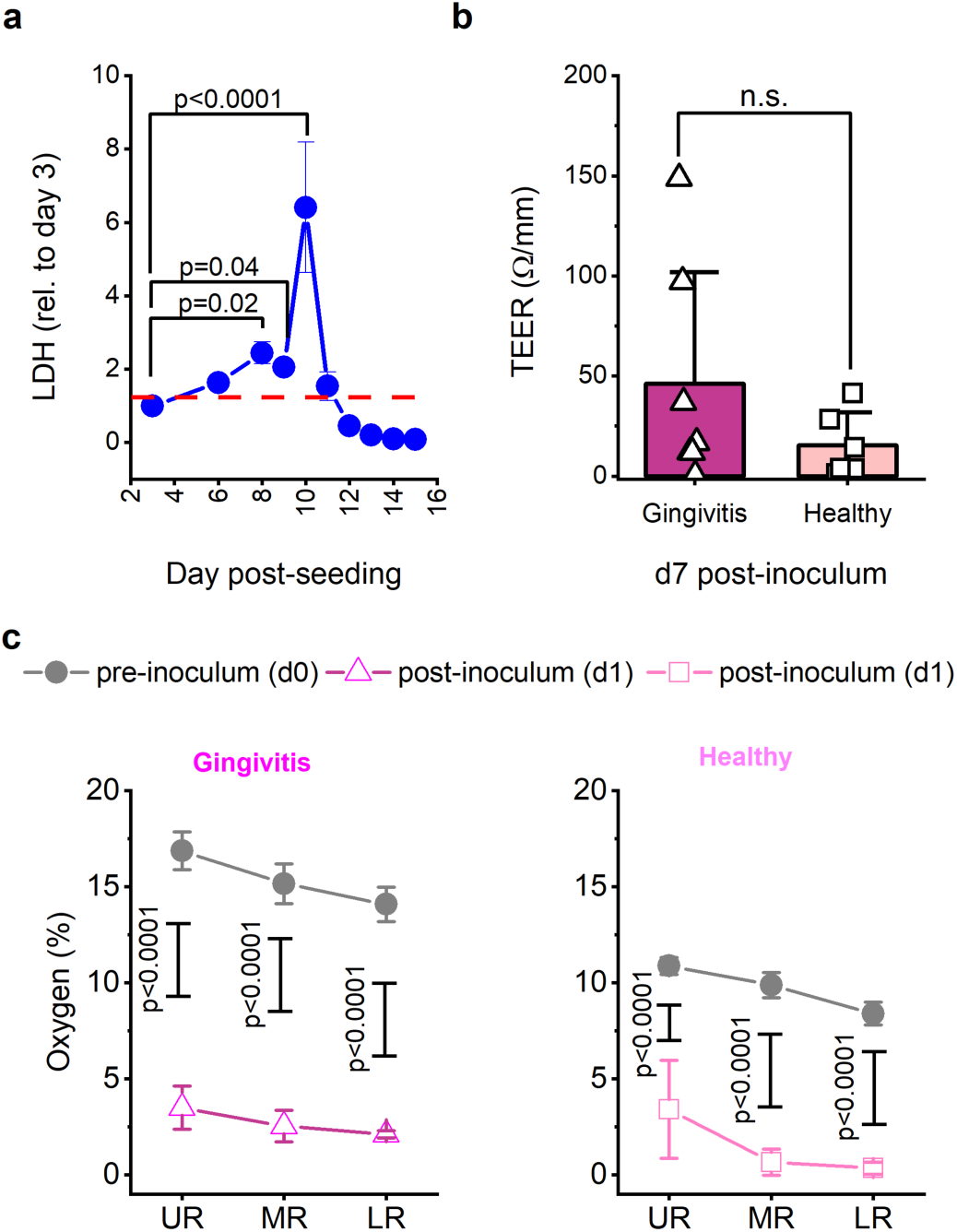
OTM remains viable and functional upon microbial colonization and distribution at the gingival sulcus. **a.** Change in LDH relative to Day 2 (dashed red line) post-gingival model fabrication and maturation. One-Way ANOVA repetitive measures with Dunnet (d2) post-hoc, p-values (<0.05) (n=5); the line plot represents the mean ± SD. **b.** TEER assessments (Ω/mm) of oral epithelium on Day 7 according to disease states (n=6 gingivitis, n=3 healthy) [37]. Two-Sample Student’s t-test, p>0.05; n.s.=not significant. **c.** Oxygen levels (%) assessed at Day 0 and Day 1 [37] at different pocket depths (Upper region (UR), medium region (MR), and lower region (LR)) within the OTM; paired t-test, p-values (<0.05).

**Supplementary Figure 2:**
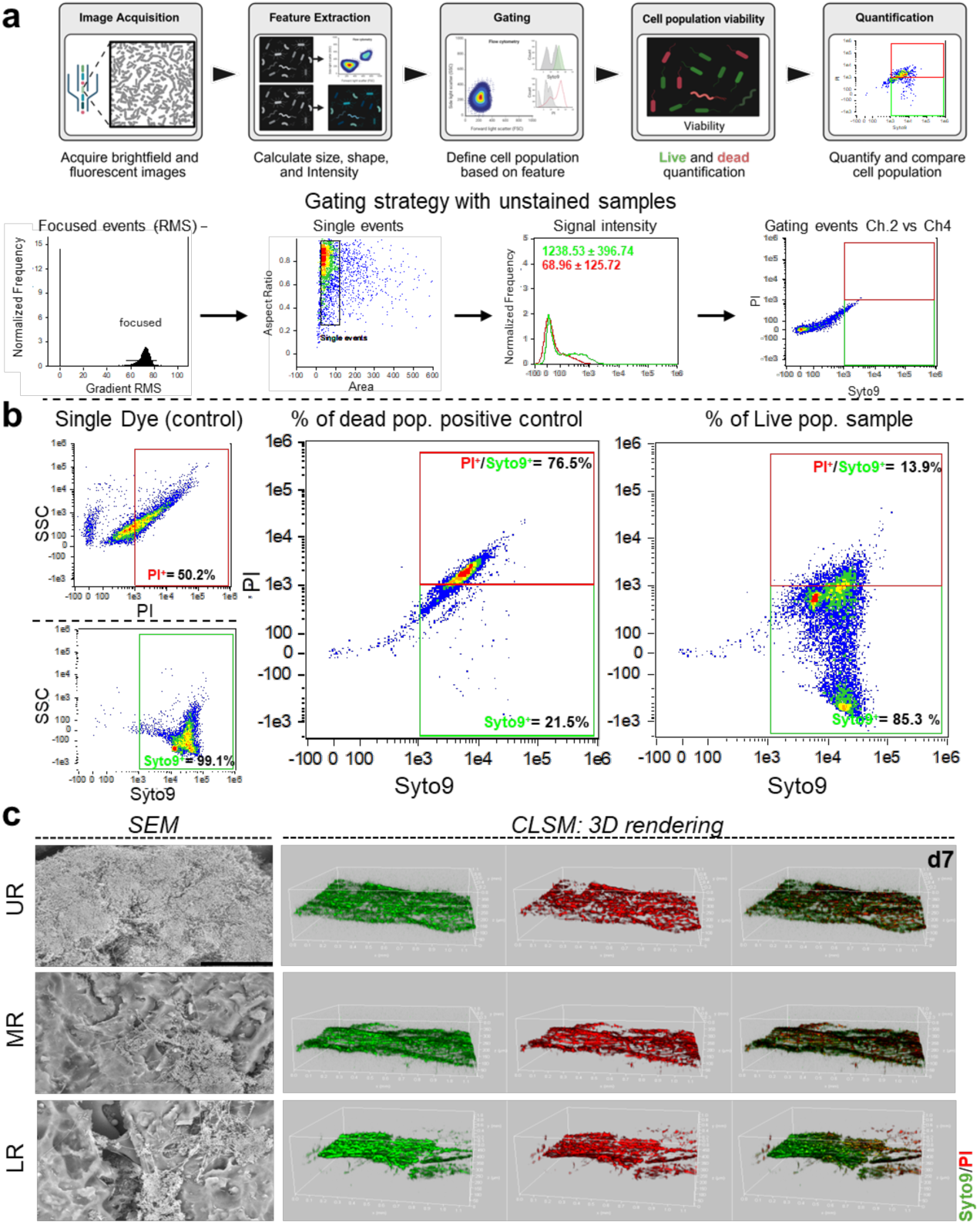
OTMs promote dysbiotic colonization of the microbiome, organization along the depth of the pocket, and long-term viability. **a-b:** Flow cytometry pipeline and viability analysis of gingivitis microbiomes isolated from the OTMs on Day 7 via density gradient centrifugation (Percoll®) technique. **a.** Upper — diagram outlining the experimental workflow for quantitative live/dead assessments; bottom — gating strategy applied for data analysis based on negative control (unstained sample); left to right — distribution of raw mean square (RMS) gradient values as a function of normalized single-event frequency for Syto9 (green) and propidium iodide (PI, red). The rectangular region highlights the events used for gating. **b.** Representative flow cytometry plots of single-dye controls (live cells - SYTO9⁺), (dead cells - PI⁺), and double-stained positive controls for dead and live populations (SYTO9⁺/PI⁺). Each plot reports the percentage of positive microbial cells for each dye. n=2. **c.** SEM (n=2) and CLSM (n=2) qualitative assessments of microbial population organization and viability within the depth of the gingival pocket on Day 7. SEM scale bar = 30 µm. Cartoon was made in Biorender.com.

**Supplementary Figure 3:**
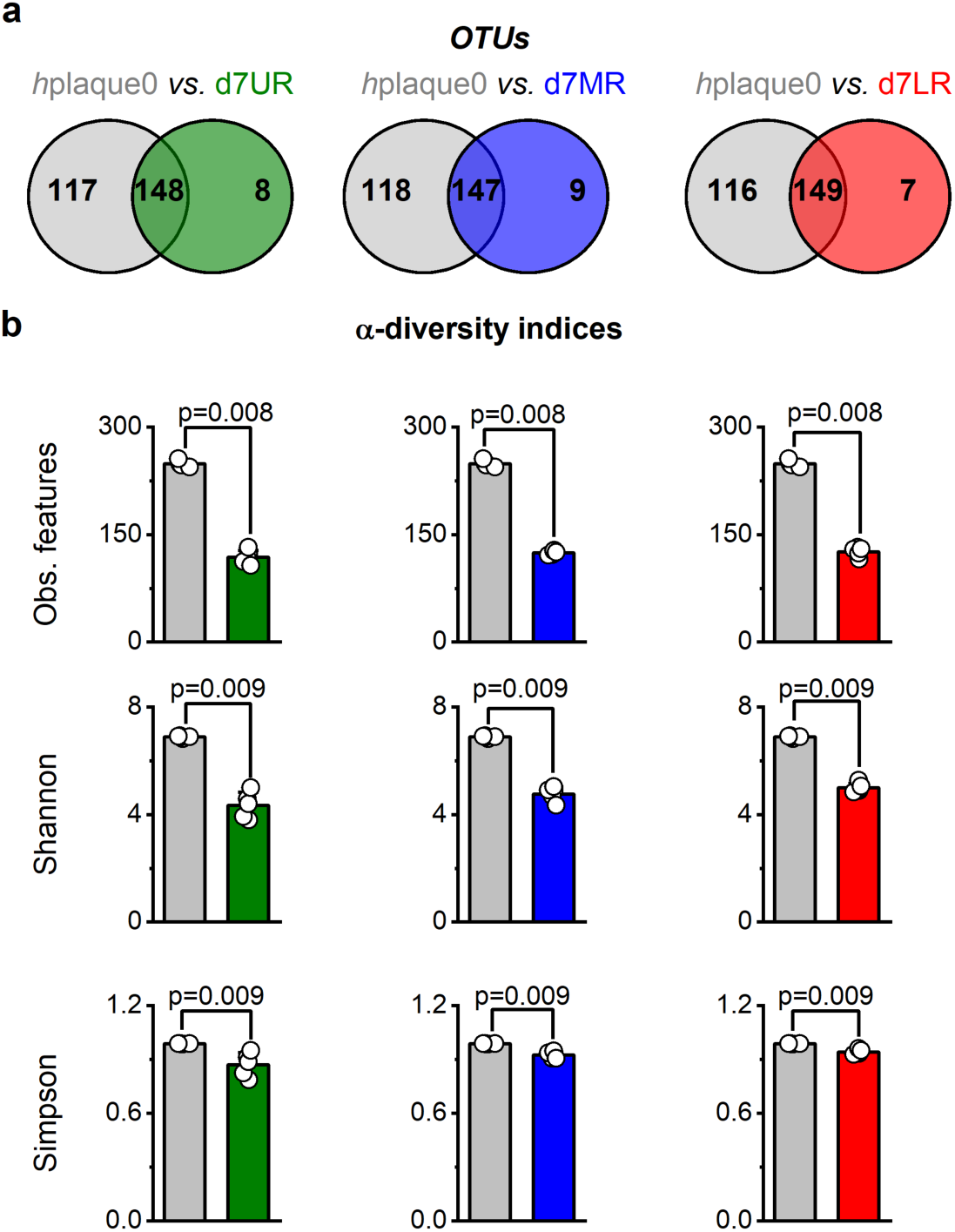
The anatomical architecture of the OTM supports the establishment of ecological niches within the pocket depth. Comparisons between gingival sulcus depths (regionality) in the OTM on d7: upper region (UR — green), medium region (MR — blue), lower region (LR — red), n=5: **a.** Venn analysis — species. **b:** α-diversity indices — Observed features, Shannon and Simpson; Kruskal–Wallis H test, p-values (<0.05) are shown in the figure.

**Supplementary Figure 4.**
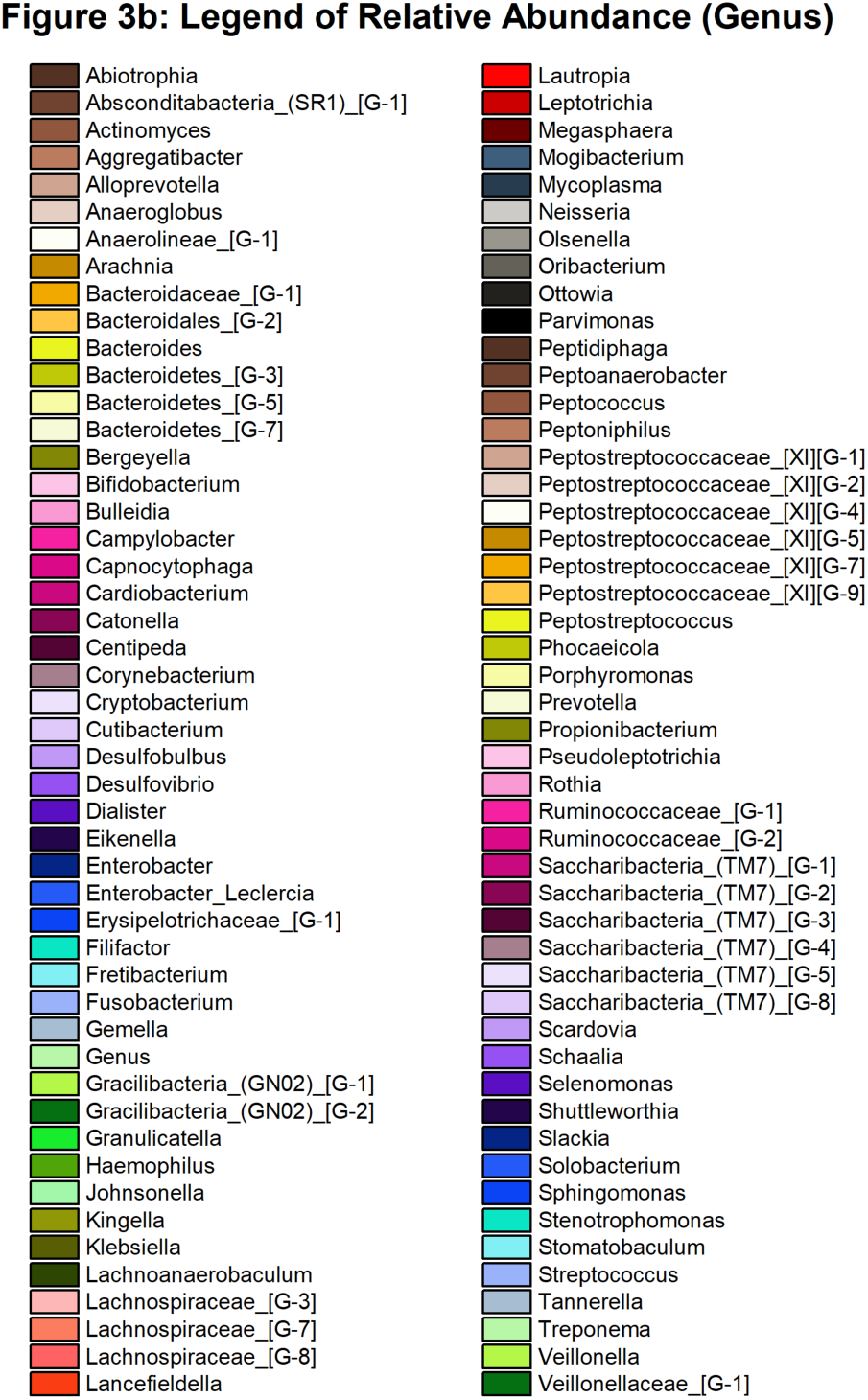
Legend of relative abundance shown in Figure 3b.

**Supplementary Figure 5.**
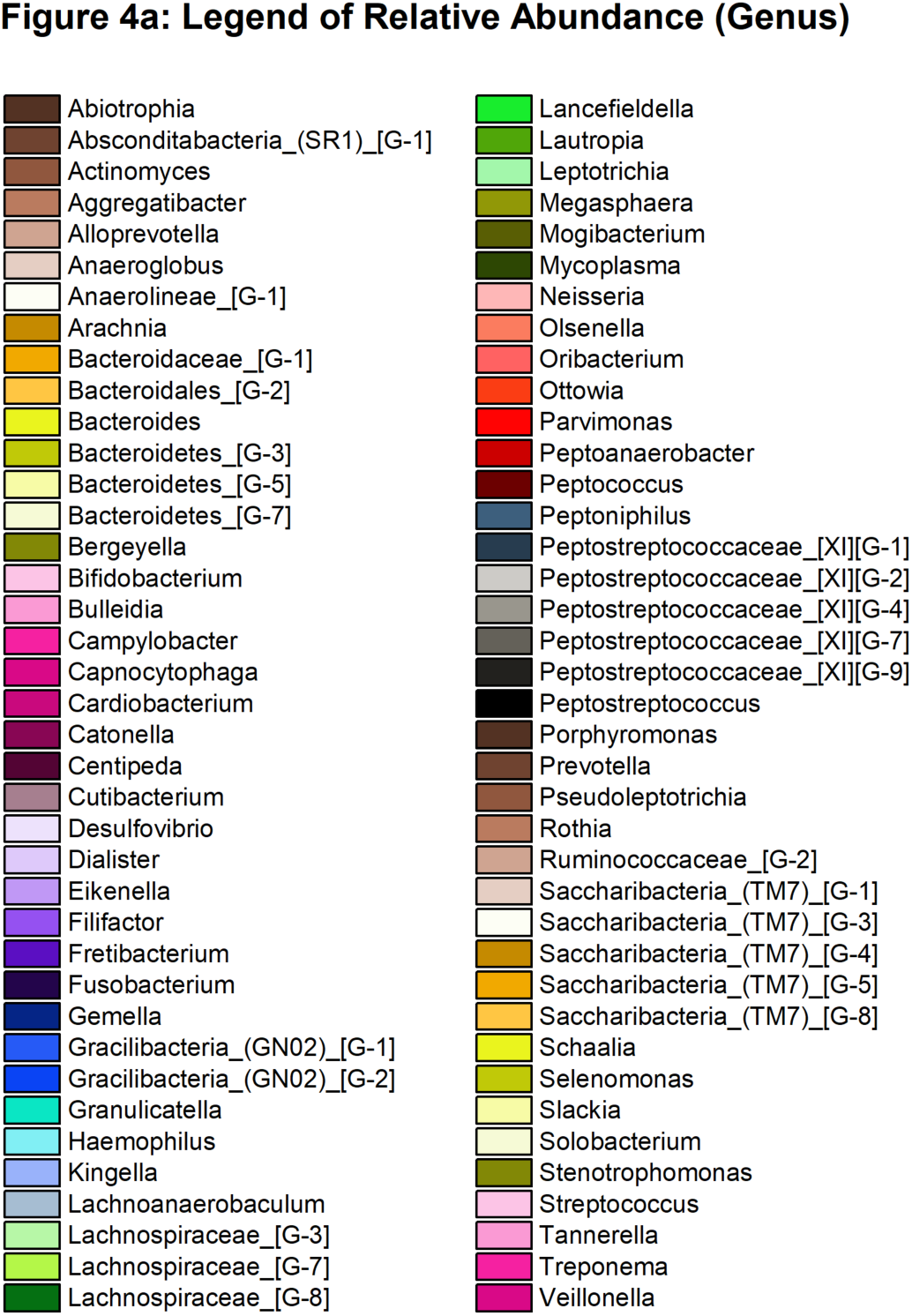
Legend of relative abundance shown in Figure 4a.

**Supplementary Figure 6:**
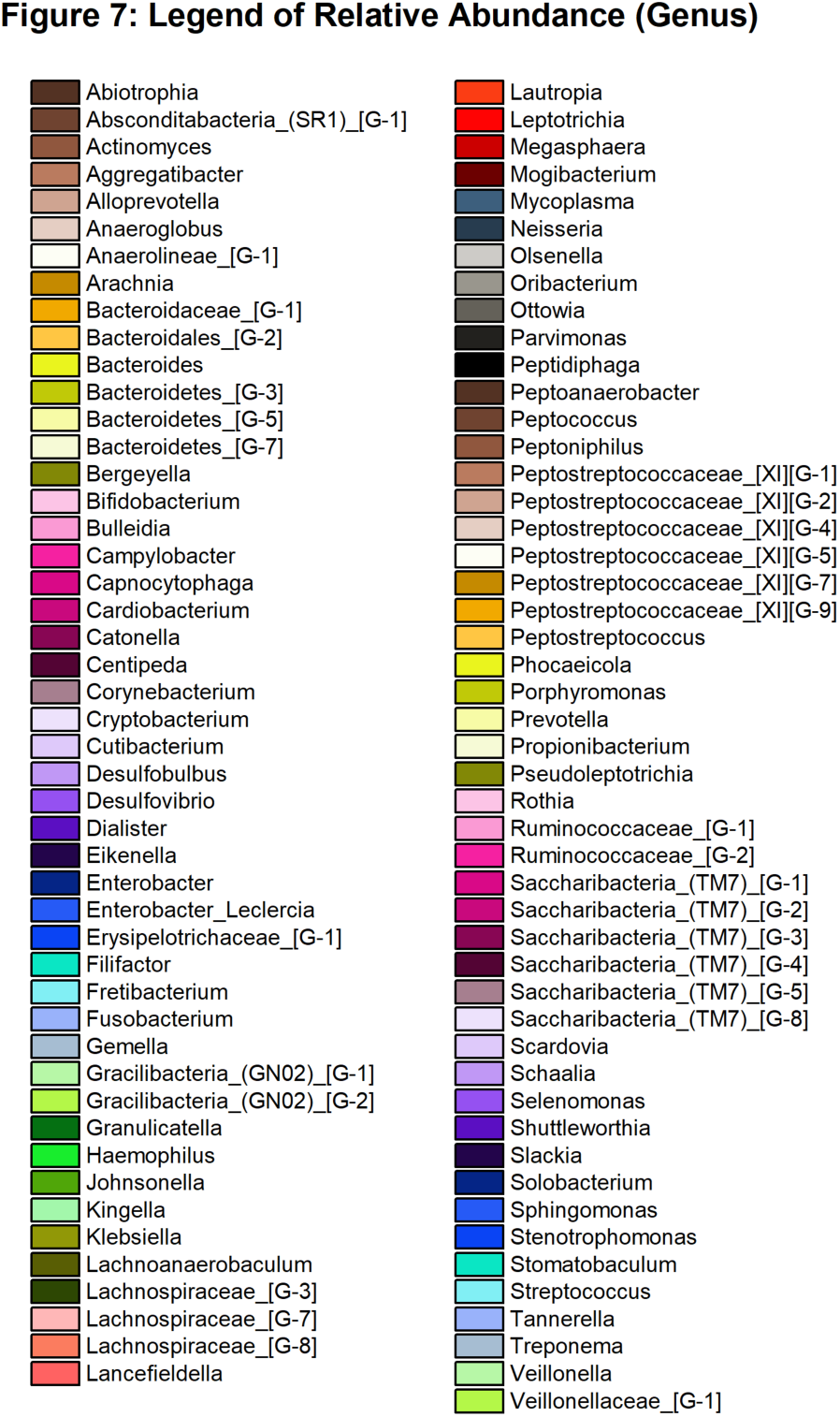
Legend of relative abundance shown in Figure 7a.

## Supplementary Tables

**Supplementary Table 1:** Enrollment of human subjects (gingivitis) in the proposed studies.

| Racial Categories | Ethnic Categories |  |  |  | Total |
| --- | --- | --- | --- | --- | --- |
|  | Not Hispanic or Latino |  | Hispanic or Latino |  |  |
|  | Female | Male | Female | Male |  |
| American Indian/ Alaska Native | 0 | 0 | 0 | 0 | 0 |
| Asian | 1 | 1 | 0 | 0 | 2 |
| Native Hawaiian or Other Pacific Islander | 0 | 0 | 0 | 0 | 0 |
| Black or African American | 2 | 2 | 0 | 0 | 4 |
| White | 5 | 5 | 2 | 2 | 14 |
| More than One Race | 0 | 0 | 0 | 0 | 0 |
| Total | 8 | 8 | 2 | 2 | 20 |

**Supplementary Table 2:** Reported p-values of the statistical analysis in Figure 6b. Two-Way repeated-measures ANOVA with Bonferroni post-hoc test; p-values (<0.05) are reported below. n.s.= not significant.

| IL-1α - OTMs inoculated with healthy subgingival microbiome |  |
| --- | --- |
| Time point | p-value |
| d1 vs. d3 | n.s. |
| d1 vs. d7 | n.s. |
| d3 vs. d7 | n.s. |
| IL-1α - OTM inoculated with gingivitis subgingival microbiome |  |
| Time point | p-value |
| d1 vs. d3 | 8.38x10<sup>-4</sup> |
| d1 vs. d7 | 2.78 x10<sup>-4</sup> |
| d3 vs. d7 | n.s. |
| IL-1α - OTMs inoculated with healthy vs. OTM inoculated with gingivitis subgingival microbiomes |  |
| Time point | p-value |
| d1 | 0.004 |
| d3 | n.s. |
| d7 | n.s. |

| **IL-1β - OTMs inoculated with healthy subgingival microbiome** |  |
| --- | --- |
| Time point | p-value |
| d1 vs. d3 | <0.0001 |
| d1 vs. d7 | <0.0001 |
| d3 vs. d7 | n.s. |
| **IL-1β - OTMs inoculated with gingivitis subgingival microbiome** |  |
| Time point | p-value |
| d1 vs. d3 | <0.0001 |
| d1 vs. d7 | <0.0001 |
| d3 vs. d7 | n.s. |
| **IL-1β - OTMs inoculated with healthy vs. OTM inoculated with gingivitis subgingival microbiomes** |  |
| Time point | p-value |
| d1 | <0.0001 |
| d3 | n.s. |
| d7 | n.s. |
| **IL-8 - OTMs inoculated with healthy subgingival microbiome** |  |
| Time point | p-value |
| d1 vs. d3 | <0.0001 |
| d1 vs. d7 | <0.0001 |
| d3 vs. d7 | n.s. |
| **IL-8 - OTMs inoculated with gingivitis subgingival microbiome** |  |
| Time point | p-value |
| d1 vs. d3 | <0.0001 |
| d1 vs. d7 | <0.0001 |
| d3 vs. d7 | n.s. |
| **IL-8 - OTMs inoculated with healthy vs. OTM inoculated with gingivitis subgingival microbiomes** |  |
| Time point | p-value |
| d1 | n.s. |
| d3 | n.s. |
| d7 | n.s. |

| MCP1 - OTMs inoculated with healthy subgingival microbiome |  |
| --- | --- |
| Time point | p-value |
| d1 vs. d3 | 0.01 |
| d1 vs. d7 | 0.01 |
| d3 vs. d7 | n.s. |
| MCP1 - OTMs inoculated with gingivitis subgingival microbiome |  |
| Time point | p-value |
| d1 vs. d3 | <0.0001 |
| d1 vs. d7 | <0.0001 |
| d3 vs. d7 | n.s. |
| MCP1- OTMs inoculated with healthy vs. OTM inoculated with gingivitis subgingival microbiomes |  |
| Time point | p-value |
| d1 | <0.0001 |
| d3 | n.s. |
| d7 | n.s. |

**Supplementary Table 3:** Reported p-values of the statistical analysis in Figure 8a. Two-Way repeated-measures ANOVA with Bonferroni post hoc test and One-Way repeated measures ANOVA with Dunnett post-hoc test (d0). n.s.= not significant.

| *OTMs inoculated with healthy subgingival microbiome* |  |  |  |  |  |  |  |  |
| --- | --- | --- | --- | --- | --- | --- | --- | --- |
| Time point | Glutamine | Glutamate | Glucose | Sodium | Calcium | Potassium | Ammonium | Lactate |
| d1 vs. d0 | n.s. | n.s. | n.s. | n.s. | n.s. | n.s. | 0.00977 | 1.15x10<sup>-5</sup> |
| d2 vs. d0 | n.s. | n.s. | n.s. | n.s. | n.s. | n.s. | 2.88x10<sup>-7</sup> | 1.54x10<sup>-5</sup> |
| d3 vs. d0 | n.s. | n.s. | n.s. | n.s. | n.s. | n.s. | 1.28x10<sup>-9</sup> | 4.49x10<sup>-8</sup> |
| d4 vs. d0 | n.s. | n.s. | n.s. | n.s. | n.s. | n.s. | 5.57x10<sup>-11</sup> | 1.88x10<sup>-8</sup> |
| d5 vs. d0 | n.s. | n.s. | n.s. | n.s. | n.s. | n.s. | 2.33x10<sup>-11</sup> | 1.88x10<sup>-8</sup> |
| d6 vs. d0 | n.s. | n.s. | n.s. | n.s. | n.s. | n.s. | 2.08x10<sup>-11</sup> | 1.07x10<sup>-7</sup> |
| d7 vs. d0 | n.s. | n.s. | n.s. | n.s. | n.s. | n.s. | 5.68x10<sup>-13</sup> | 1.07x10<sup>-7</sup> |
| *OTMs inoculated with gingivitis subgingival microbiome* |  |  |  |  |  |  |  |  |
| Time point | Glutamine | Glutamate | Glucose | Sodium | Calcium | Potassium | Ammonium | Lactate |
| d1 vs. d0 | n.s. | n.s. | n.s. | 0.013 | 0.02 | n.s. | 4.78x10<sup>-8</sup> | 2.54x10<sup>-10</sup> |
| d2 vs. d0 | 1.01x10<sup>-14</sup> | 0.01 | n.s. | n.s. | n.s. | n.s. | 1.12x10<sup>-9</sup> | 1.24x10<sup>-10</sup> |
| d3 vs. d0 | <0.001 | 0.01 | n.s. | n.s. | n.s. | n.s. | <0.001 | 1.32x10<sup>-10</sup> |
| d4 vs. d0 | <0.001 | 0.01 | n.s. | n.s. | n.s. | n.s. | <0.001 | 1.51x10<sup>-10</sup> |
| d5 vs. d0 | <0.001 | n.s. | n.s. | n.s. | n.s. | 0.004 | <0.001 | 0.003 |
| d6 vs. d0 | <0.001 | 0.01 | n.s. | n.s. | n.s. | n.s. | <0.001 | 1.51x10<sup>-10</sup> |
| d7 vs. d0 | <0.001 | n.s. | n.s. | 0.0365 | n.s. | n.s. | <0.001 | 2.09x10<sup>-10</sup> |
| *OTMs inoculated with healthy vs. OTM inoculated with gingivitis subgingival microbiomes* |  |  |  |  |  |  |  |  |
| Time point | Glutamine | Glutamate | Glucose | Sodium | Calcium | Potassium | Ammonium | Lactate |
| d0 | n.s. | n.s. | 0.01 | n.s. | n.s. | n.s. | n.s. | 0.004 |
| d1 | n.s. | n.s. | 1.47x10<sup>-4</sup> | n.s. | n.s. | n.s. | 0.007 | n.s. |
| d2 | 2.69x10<sup>-8</sup> | n.s. | 1.52x10<sup>-4</sup> | n.s. | n.s. | n.s. | 0.008 | n.s. |
| d3 | 5.86x10<sup>-37</sup> | n.s. | 1.21x10<sup>-4</sup> | 0.007 | n.s. | n.s. | 3.30x10<sup>-12</sup> | n.s. |
| d4 | 3.68x10<sup>-36</sup> | n.s. | 4.03x10<sup>-4</sup> | n.s. | n.s. | n.s. | 7.79x10<sup>-13</sup> | n.s. |
| d5 | 2.14x10<sup>-36</sup> | n.s. | n.s. | 0.024 | 1.23x10<sup>-10</sup> | 5.05x10<sup>-4</sup> | 1.37x10<sup>-8</sup> | n.s. |
| d6 | 7.50x10<sup>-37</sup> | n.s. | 3.88x10<sup>-5</sup> | n.s. | 0.02 | n.s. | 1.04x10<sup>-14</sup> | n.s. |
| d7 | 1.69x10<sup>-33</sup> | n.s. | 6.05x10<sup>-4</sup> | n.s. | n.s. | n.s. | 1.52x10<sup>-11</sup> | n.s. |

**Supplementary Table 4:** Reported p-values of the statistical analysis in Figure 8b. One-Way repeated measures ANOVA with Dunnett post-hoc test (d0). n.s.= not significant.

| pH recorded in OTMs inoculated with healthy subgingival microbiome |  |
| --- | --- |
| Time point | p-value |
| d1 vs. d0 | <0.0001 |
| d2 vs. d0 | <0.0001 |
| d3 vs. d0 | <0.0001 |
| d4 vs. d0 | <0.0001 |
| d5 vs. d0 | <0.0001 |
| d6 vs. d0 | <0.0001 |
| d7 vs. d0 | <0.0001 |
| pH recorded in OTMs inoculated with gingivitis subgingival microbiome |  |
| Time point | p-value |
| d1 vs. d0 | <0.0001 |
| d2 vs. d0 | 0.001 |
| d3 vs. d0 | 0.001 |
| d4 vs. d0 | n.s. |
| d5 vs. d0 | <0.0001 |
| d6 vs. d0 | 0.004 |
| d7 vs. d0 | <0.0001 |

