## Supplementary Information_Community ID for "Profiling Gingival Inflammation in a 3D Oral Tissue Model Reveals Early Features of Disease Progression"

| **TaxonID** | **Species** | **Module ID** | **Presence** |
| --- | --- | --- | --- |
| SP146 | Peptoanaerobacter [Eubacterium] yurii | 1 | Day7 |
| SP144 | Peptostreptococcaceae_[XI][G-1] [Eubacterium]_infirmum | 1 | Both |
| SP69 | Filifactor alocis | 1 | Both |
| SP43 | Streptococcus anginosus | 1 | Both |
| SP114 | Veillonella atypica | 1 | Day7 |
| SP128 | Peptostreptococcaceae_[XI][G-7] bacterium_HMT_081 | 1 | Both |
| SP120 | Ruminococcaceae_[G-2] bacterium_HMT_085 | 1 | Both |
| SP173 | Peptostreptococcaceae_[XI][G-2] bacterium_HMT_091 | 1 | Both |
| SP185 | Lachnospiraceae_[G-7] bacterium_HMT_163 | 1 | Both |
| SP364 | Bacteroidaceae_[G-1] bacterium_HMT_272 | 1 | Both |
| SP155 | Bacteroidetes_[G-3] bacterium_HMT_280 | 1 | Both |
| SP15 | Absconditabacteria_(SR1)_[G-1] bacterium_HMT_345 | 1 | Both |
| SPN359 | Saccharibacteria_(TM7)_[G-3] bacterium_HMT_351_nov_92.871% | 1 | Day7 |
| SP216 | Saccharibacteria_(TM7)_[G-4] bacterium_HMT_355 | 1 | Both |
| SP296 | Peptostreptococcaceae_[XI][G-4] bacterium_HMT_369 | 1 | Both |
| SP389 | Bacteroidetes_[G-3] bacterium_HMT_436 | 1 | Day7 |
| SP21 | Lachnospiraceae_[G-8] bacterium_HMT_500 | 1 | Both |
| SP131 | Gracilibacteria_(GN02)_[G-2] bacterium_HMT_873 | 1 | Day7 |
| SP375 | Prevotella baroniae | 1 | Both |
| SP163 | Prevotella buccae | 1 | Both |
| SP354 | Selenomonas dianae | 1 | Day7 |
| SP60 | Veillonella dispar | 1 | Both |
| SPP25 | Veillonella dispar_parvula | 1 | Both |
| SP42 | Neisseria elongata | 1 | Both |
| SP32 | Porphyromonas endodontalis | 1 | Both |
| SP250 | Slackia exigua | 1 | Both |
| SP294 | Bulleidia extructa | 1 | Both |
| SP80 | Neisseria flava | 1 | Both |
| SP164 | Neisseria flavescens | 1 | Day7 |
| SPN427 | Tannerella forsythia_nov_97.308% | 1 | Both |
| SP380 | Porphyromonas gingivalis | 1 | Both |
| SP23 | Prevotella intermedia | 1 | Both |
| SP301 | Peptoniphilus lacrimalis | 1 | Both |
| SPN40 | Desulfovibrio legallii_nov_94.343% | 1 | Both |
| SP2 | Stenotrophomonas maltophilia | 1 | Day7 |
| SP27 | Solobacterium moorei | 1 | Both |
| SP26 | Catonella morbi | 1 | Both |
| SP282 | Neisseria mucosa | 1 | Both |
| SPP13 | Fusobacterium nucleatum_nucleatum_subsp._animalis | 1 | Both |
| SP209 | Fusobacterium nucleatum_subsp._animalis | 1 | Both |
| SP153 | Fusobacterium nucleatum_subsp._vincentii | 1 | Both |
| SP112 | Streptococcus oralis | 1 | Both |
| SP243 | Prevotella oralis | 1 | Both |
| SP33 | Streptococcus oralis_subsp._tigurinus_clade_070 | 1 | Both |
| SP121 | Streptococcus oralis_subsp._tigurinus_clade_071 | 1 | Day7 |
| SP275 | Lancefieldella parvula | 1 | Both |
| SP38 | Veillonella parvula | 1 | Both |
| SP345 | Dialister pneumosintes | 1 | Both |
| SP162 | Veillonella rogosae | 1 | Both |
| SP94 | Prevotella saccharolytica | 1 | Both |
| SP37 | Streptococcus salivarius | 1 | Both |
| SP252 | Aggregatibacter segnis | 1 | Both |
| SP55 | Campylobacter showae | 1 | Both |
| SP36 | Neisseria sicca | 1 | Both |
| SP210 | Selenomonas sp._HMT_138 | 1 | Both |
| SPN371 | Prevotella sp._HMT_305_nov_93.661% | 1 | Both |
| SP302 | Parvimonas sp._HMT_393 | 1 | Both |
| SP127 | Aggregatibacter sp._HMT_512 | 1 | Both |
| SP278 | Haemophilus sputorum | 1 | Both |
| SP110 | Peptostreptococcus stomatis | 1 | Both |
| SP106 | Neisseria subflava | 1 | Both |
| SPN105 | Bergeyella zoohelcum_nov_92.593% | 1 | Both |
| SP260 | Peptostreptococcaceae_[XI][G-9] [Eubacterium]_brachy | 2 | Both |
| SP159 | Peptostreptococcaceae_[XI][G-5] [Eubacterium]_saphenum | 2 | Day0 |
| SP108 | Peptostreptococcaceae_[XI][G-7] [Eubacterium]_yurii_subsps._yurii_&_margaretiae | 2 | Both |
| SP48 | Granulicatella adiacens | 2 | Both |
| SP208 | Rothia aeria | 2 | Both |
| SP150 | Aggregatibacter aphrophilus | 2 | Both |
| SP125 | Selenomonas artemidis | 2 | Day0 |
| SP365 | Neisseria bacilliformis | 2 | Both |
| SP286 | Ruminococcaceae_[G-1] bacterium_HMT_075 | 2 | Day0 |
| SP77 | Lachnospiraceae_[G-3] bacterium_HMT_100 | 2 | Both |
| SP231 | Peptostreptococcaceae_[XI][G-4] bacterium_HMT_103 | 2 | Day0 |
| SP168 | Veillonellaceae_[G-1] bacterium_HMT_155 | 2 | Day0 |
| SP35 | Bacteroidales_[G-2] bacterium_HMT_274 | 2 | Both |
| SP47 | Saccharibacteria_(TM7)_[G-1] bacterium_HMT_346 | 2 | Both |
| SP258 | Saccharibacteria_(TM7)_[G-1] bacterium_HMT_348 | 2 | Day0 |
| SP109 | Saccharibacteria_(TM7)_[G-1] bacterium_HMT_349 | 2 | Day0 |
| SP175 | Saccharibacteria_(TM7)_[G-2] bacterium_HMT_350 | 2 | Day0 |
| SP182 | Saccharibacteria_(TM7)_[G-5] bacterium_HMT_356 | 2 | Both |
| SP220 | Anaerolineae_[G-1] bacterium_HMT_439 | 2 | Both |
| SP273 | Bacteroidetes_[G-5] bacterium_HMT_511 | 2 | Both |
| SP274 | Saccharibacteria_(TM7)_[G-1] bacterium_HMT_869 | 2 | Day0 |
| SP41 | Gracilibacteria_(GN02)_[G-1] bacterium_HMT_872 | 2 | Both |
| SP304 | Absconditabacteria_(SR1)_[G-1] bacterium_HMT_874 | 2 | Day0 |
| SP74 | Erysipelotrichaceae_[G-1] bacterium_HMT_905 | 2 | Day0 |
| SP215 | Saccharibacteria_(TM7)_[G-1] bacterium_HMT_957 | 2 | Both |
| SP20 | Leptotrichia buccalis | 2 | Day0 |
| SP1 | Porphyromonas catoniae | 2 | Both |
| SP81 | Streptococcus chosunense | 2 | Day0 |
| SP249 | Campylobacter concisus | 2 | Day0 |
| SP200 | Streptococcus constellatus | 2 | Both |
| SP9 | Eikenella corrodens | 2 | Both |
| SP213 | Cryptobacterium curtum | 2 | Day0 |
| SP103 | Abiotrophia defectiva | 2 | Both |
| SP367 | Kingella denitrificans | 2 | Day0 |
| SP157 | Treponema denticola | 2 | Both |
| SP401 | Bifidobacterium dentium | 2 | Both |
| SP78 | Rothia dentocariosa | 2 | Both |
| SP66 | Mogibacterium diversum | 2 | Day0 |
| SP211 | Corynebacterium durum | 2 | Day0 |
| SPN18 | Corynebacterium durum_nov_97.787% | 2 | Day0 |
| SP140 | Capnocytophaga endodontalis | 2 | Both |
| SP134 | Fretibacterium fastidiosum | 2 | Both |
| SP371 | Mycoplasma faucium | 2 | Both |
| SP184 | Tannerella forsythia | 2 | Both |
| SP295 | Anaeroglobus geminatus | 2 | Day0 |
| SP14 | Capnocytophaga gingivalis | 2 | Both |
| SP262 | Lachnoanaerobaculum gingivalis | 2 | Both |
| SPN339 | Capnocytophaga gingivalis_nov_90.756% | 2 | Both |
| SPN451 | Capnocytophaga gingivalis_nov_96.218% | 2 | Day0 |
| SPP16 | Lachnoanaerobaculum gingivalis_umeaense | 2 | Day0 |
| SP196 | Peptidiphaga gingivicola | 2 | Day0 |
| SPN213 | Peptidiphaga gingivicola_nov_96.920% | 2 | Day0 |
| SP290 | Pseudoleptotrichia goodfellowii | 2 | Both |
| SP179 | Campylobacter gracilis | 2 | Day0 |
| SP266 | Capnocytophaga granulosa | 2 | Both |
| SP124 | Gemella haemolysans | 2 | Both |
| SP132 | Capnocytophaga haemolytica | 2 | Day0 |
| SP67 | Leptotrichia hofstadii | 2 | Day0 |
| SP101 | Cardiobacterium hominis | 2 | Both |
| SP130 | Leptotrichia hongkongensis | 2 | Day0 |
| SP193 | Fusobacterium hwasookii | 2 | Both |
| SP419 | Johnsonella ignava | 2 | Day0 |
| SPP1 | Streptococcus infantis_infantis_clade_638 | 2 | Day0 |
| SP54 | Streptococcus intermedius | 2 | Both |
| SP187 | Dialister invisus | 2 | Both |
| SP13 | Capnocytophaga leadbetteri | 2 | Both |
| SP305 | Treponema lecithinolyticum | 2 | Day0 |
| SP90 | Prevotella loescheii | 2 | Both |
| SP181 | Stomatobaculum longum | 2 | Day0 |
| SP87 | Prevotella maculosa | 2 | Both |
| SP177 | Actinomyces massiliensis | 2 | Day0 |
| SP100 | Corynebacterium matruchotii | 2 | Day0 |
| SPN404 | Corynebacterium matruchotii_nov_97.951% | 2 | Day0 |
| SP50 | Prevotella melaninogenica | 2 | Day0 |
| SP362 | Schaalia meyeri | 2 | Day0 |
| SP330 | Prevotella micans | 2 | Day0 |
| SP12 | Parvimonas micra | 2 | Both |
| SP267 | Megasphaera micronuciformis | 2 | Day0 |
| SP75 | Lautropia mirabilis | 2 | Both |
| SP97 | Streptococcus mitis | 2 | Day0 |
| SP248 | Streptococcus mutans | 2 | Day0 |
| SP61 | Actinomyces naeslundii | 2 | Day0 |
| SP160 | Prevotella nigrescens | 2 | Both |
| SP63 | Selenomonas noxia | 2 | Day0 |
| SPN49 | Selenomonas noxia_nov_97.446% | 2 | Day0 |
| SP62 | Fusobacterium nucleatum | 2 | Both |
| SP165 | Capnocytophaga ochracea | 2 | Both |
| SP333 | Schaalia odontolytica | 2 | Both |
| SP40 | Lachnoanaerobaculum orale | 2 | Both |
| SP237 | Kingella oralis | 2 | Both |
| SP245 | Neisseria oralis | 2 | Both |
| SP151 | Prevotella oulorum | 2 | Both |
| SP24 | Haemophilus parahaemolyticus | 2 | Both |
| SP53 | Haemophilus parainfluenzae | 2 | Both |
| SP46 | Porphyromonas pasteri | 2 | Both |
| SP195 | Fusobacterium periodonticum | 2 | Both |
| SP65 | Centipeda periodontii | 2 | Both |
| SPN383 | Centipeda periodontii_nov_97.436% | 2 | Both |
| SP99 | Prevotella pleuritidis | 2 | Day0 |
| SP76 | Arachnia propionica | 2 | Both |
| SP310 | Alloprevotella rava | 2 | Day0 |
| SP293 | Campylobacter rectus | 2 | Both |
| SP88 | Lancefieldella rimae | 2 | Day0 |
| SP337 | Arachnia rubra | 2 | Day0 |
| SP166 | Lachnoanaerobaculum saburreum | 2 | Both |
| SP170 | Prevotella salivae | 2 | Day0 |
| SP308 | Mycoplasma salivarium | 2 | Both |
| SP136 | Shuttleworthia satelles | 2 | Day0 |
| SP116 | Tannerella serpentiformis | 2 | Day0 |
| SPP4 | Tannerella serpentiformis_sp._HMT_286 | 2 | Day0 |
| SP253 | Leptotrichia shahii | 2 | Day0 |
| SP261 | Oribacterium sinus | 2 | Day0 |
| SP331 | Desulfobulbus sp._HMT_041 | 2 | Day0 |
| SP138 | Oribacterium sp._HMT_078 | 2 | Both |
| SP259 | Lachnoanaerobaculum sp._HMT_083 | 2 | Day0 |
| SP84 | Parvimonas sp._HMT_110 | 2 | Day0 |
| SP268 | Megasphaera sp._HMT_123 | 2 | Both |
| SP232 | Selenomonas sp._HMT_137 | 2 | Day0 |
| SPN267 | Selenomonas sp._HMT_137_nov_97.137% | 2 | Day0 |
| SP28 | Peptococcus sp._HMT_167 | 2 | Both |
| SP73 | Actinomyces sp._HMT_169 | 2 | Both |
| SP107 | Actinomyces sp._HMT_170 | 2 | Day0 |
| SP93 | Actinomyces sp._HMT_171 | 2 | Day0 |
| SPN416 | Actinomyces sp._HMT_171_nov_96.929% | 2 | Day0 |
| SP34 | Actinomyces sp._HMT_175 | 2 | Both |
| SP137 | Schaalia sp._HMT_180 | 2 | Both |
| SP98 | Peptidiphaga sp._HMT_183 | 2 | Day0 |
| SP123 | Fusobacterium sp._HMT_203 | 2 | Both |
| SP228 | Fusobacterium sp._HMT_204 | 2 | Day0 |
| SP71 | Leptotrichia sp._HMT_212 | 2 | Both |
| SP191 | Leptotrichia sp._HMT_215 | 2 | Day0 |
| SP240 | Leptotrichia sp._HMT_219 | 2 | Both |
| SP379 | Leptotrichia sp._HMT_223 | 2 | Day0 |
| SP113 | Leptotrichia sp._HMT_225 | 2 | Day0 |
| SP272 | Treponema sp._HMT_231 | 2 | Both |
| SP192 | Treponema sp._HMT_237 | 2 | Day0 |
| SP328 | Treponema sp._HMT_247 | 2 | Day0 |
| SP115 | Treponema sp._HMT_270 | 2 | Day0 |
| SP52 | Porphyromonas sp._HMT_275 | 2 | Day0 |
| SP229 | Porphyromonas sp._HMT_284 | 2 | Both |
| SP11 | Tannerella sp._HMT_286 | 2 | Day0 |
| SP25 | Prevotella sp._HMT_300 | 2 | Both |
| SP219 | Prevotella sp._HMT_301 | 2 | Day0 |
| SP201 | Prevotella sp._HMT_314 | 2 | Day0 |
| SP95 | Prevotella sp._HMT_315 | 2 | Both |
| SP148 | Prevotella sp._HMT_317 | 2 | Both |
| SP343 | Capnocytophaga sp._HMT_326 | 2 | Day0 |
| SP171 | Capnocytophaga sp._HMT_332 | 2 | Day0 |
| SP139 | Capnocytophaga sp._HMT_336 | 2 | Both |
| SP257 | Capnocytophaga sp._HMT_338 | 2 | Day0 |
| SP256 | Fretibacterium sp._HMT_359 | 2 | Day0 |
| SP5 | Fretibacterium sp._HMT_360 | 2 | Both |
| SP118 | Fusobacterium sp._HMT_370 | 2 | Both |
| SP4 | Leptotrichia sp._HMT_392 | 2 | Day0 |
| SP376 | Selenomonas sp._HMT_442 | 2 | Both |
| SP254 | Actinomyces sp._HMT_448 | 2 | Day0 |
| SP72 | Aggregatibacter sp._HMT_458 | 2 | Day0 |
| SP234 | Prevotella sp._HMT_472 | 2 | Both |
| SP224 | Alloprevotella sp._HMT_473 | 2 | Both |
| SP147 | Leptotrichia sp._HMT_498 | 2 | Both |
| SP119 | Olsenella sp._HMT_807 | 2 | Day0 |
| SP83 | Tannerella sp._HMT_808 | 2 | Both |
| SP29 | Capnocytophaga sp._HMT_864 | 2 | Day0 |
| SP271 | Schaalia sp._HMT_877 | 2 | Day0 |
| SP8 | Ottowia sp._HMT_894 | 2 | Both |
| SP205 | Aggregatibacter sp._HMT_898 | 2 | Both |
| SP117 | Capnocytophaga sp._HMT_902 | 2 | Both |
| SP226 | Leptotrichia sp._HMT_909 | 2 | Both |
| SP22 | Tannerella sp._HMT_916 | 2 | Both |
| SP143 | Selenomonas sp._HMT_919 | 2 | Both |
| SPN439 | Selenomonas sp._HMT_936_nov_97.030% | 2 | Both |
| SP57 | Selenomonas sputigena | 2 | Day0 |
| SP7 | Capnocytophaga sputigena | 2 | Both |
| SP59 | Alloprevotella tannerae | 2 | Both |
| SP126 | Mogibacterium timidum | 2 | Day0 |
| SP156 | Leptotrichia trevisanii | 2 | Day0 |
| SP307 | Olsenella uli | 2 | Both |
| SP263 | Lachnoanaerobaculum umeaense | 2 | Both |
| SP111 | Cardiobacterium valvarum | 2 | Day0 |
| SP223 | Prevotella veroralis | 2 | Day0 |
| SP276 | Treponema vincentii | 2 | Both |
| SP188 | Leptotrichia wadei | 2 | Both |
| SP227 | Scardovia wiggsiae | 2 | Day0 |
| SP381 | Cutibacterium acnes | 3 | Both |
| SPP14 | Peptostreptococcaceae_[XI][G-4] bacterium_HMT_103_bacterium_HMT_369 | 3 | Both |
| SP178 | Saccharibacteria_(TM7)_[G-3] bacterium_HMT_351 | 3 | Both |
| SP167 | Saccharibacteria_(TM7)_[G-8] bacterium_HMT_955 | 3 | Both |
| SP89 | Streptococcus cristatus | 3 | Both |
| SP363 | Streptococcus cristatus_clade_578 | 3 | Both |
| SP18 | Prevotella denticola | 3 | Both |
| SP3 | Streptococcus gordonii | 3 | Both |
| SP30 | Selenomonas infelix | 3 | Both |
| SP58 | Treponema maltophilum | 3 | Both |
| SP135 | Prevotella marshii | 3 | Both |
| SP207 | Gemella morbillorum | 3 | Both |
| SP129 | Streptococcus oralis_subsp._dentisani_clade_058 | 3 | Both |
| SP10 | Prevotella oris | 3 | Both |
| SP31 | Streptococcus sanguinis | 3 | Both |
| SPN146 | Streptococcus sanguinis_nov_97.782% | 3 | Both |
| SP39 | Treponema socranskii | 3 | Both |
| SP102 | Streptococcus sp._HMT_056 | 3 | Both |
| SP91 | Streptococcus sp._HMT_064 | 3 | Both |
| SP239 | Selenomonas sp._HMT_146 | 3 | Both |
| SPP9 | Prevotella sp._HMT_292_sp._HMT_300 | 3 | Both |
| SP344 | Prevotella sp._HMT_304 | 3 | Both |
| SP64 | Bergeyella sp._HMT_322 | 3 | Both |
| SP298 | Streptococcus sp._HMT_423 | 3 | Both |
| SP189 | Selenomonas sp._HMT_481 | 3 | Both |
| SP68 | Selenomonas sp._HMT_892 | 3 | Both |
| SPN28 | Selenomonas sp._HMT_892_nov_97.041% | 3 | Both |
| SP56 | Bergeyella sp._HMT_900 | 3 | Both |
| SP236 | Actinomyces timonensis | 3 | Both |
